# Phenotypic and clonal stability of antigen-inexperienced memory-like T cells across the genetic background, hygienic status, and aging

**DOI:** 10.1101/2020.09.01.277004

**Authors:** Alena Moudra, Veronika Niederlova, Jiri Novotny, Lucie Schmiedova, Jan Kubovciak, Tereza Matejkova, Ales Drobek, Michaela Pribikova, Romana Stopkova, Dagmar Cizkova, Ales Neuwirth, Juraj Michalik, Katerina Krizova, Tomas Hudcovic, Michal Kolar, Hana Kozakova, Jakub Kreisinger, Pavel Stopka, Ondrej Stepanek

## Abstract

Antigen-inexperienced memory-like T (AIMT) cells are functionally unique T cells representing one of the two largest subsets of murine CD8^+^ T cells. However, differences between laboratory inbred strains, insufficient data from germ-free mice, a complete lack of data from feral mice, and unclear relationship between AIMT cells formation during aging represent major barriers for better understanding of their biology. We performed a thorough characterization of AIMT cells from mice of different genetic background, age, and hygienic status by flow cytometry and multi-omics approaches including analyses of gene expression, TCR repertoire, and microbial colonization. Our data showed that AIMT cells are steadily present in mice independently of their genetic background and hygienic status. Despite differences in their gene expression profiles, young and aged AIMT cells originate from identical clones. We identified that CD122 discriminates two major subsets of AIMT cells in a strain-independent manner. Whereas thymic CD122^LOW^ AIMT cells (innate memory) prevail only in young animals with high thymic IL-4 production, peripheral CD122^HIGH^ AIMT cells (virtual memory) dominate in aged mice. Co-housing with feral mice changed the bacterial colonization of laboratory strains, but had only minimal effects on the CD8^+^ T-cell compartment including AIMT cells.

## Introduction

AIMT cells are divided into two subsets based on the site of their origin. Innate T cells originate in the thymus, whereas virtual memory T cells are formed in the peripheral lymphoid organs [1, 2]. In BALB/c mice and several gene knock-outs on C57BL/6 background, innate memory cells are formed in the thymus via IL-4 produced by invariant natural killer T (iNKT) cells [3–5]. Wild-type C57BL/6 mice have low numbers of IL-4-producing iNKT cells in the thymus and form virtual memory T cells in a process dependent on IL-15 presentation by peripheral CD8α DCs [2, 5–7]. Recently, it has been observed that mature CD8^+^ single positive thymic precursors of virtual memory T cells upregulate Eomes [8], a transcription factor required for their existence [7], suggesting that the differentiation process of virtual memory cells might be initiated already in the thymus and accomplished in the periphery [9]. This scenario is consistent with the observation that thymocytes developing in a neonatal thymus are more predisposed towards AIMT cell formation than thymocytes developing in an adult thymus of C57BL/6 mice [10].

In addition to these major routes, lymphopenia induces homeostatic proliferation coupled with the differentiation of peripheral naïve CD8^+^ T cells into AIMT cells, probably via IL-7 and/or IL-15 in C57BL/6 mice [11, 12]. Moreover, excess of IL-4, IL-7, or IL-15 induces the formation and/or expansion of AIMT cells [6, 13, 14]. Given the fact that these three cytokines signal through the common γ-chain [15], it is not surprising that all of them induce AIMT cells formation. Although it has been proposed that some genes such as *Tbx21/T-bet* or *Klrk1*/NKG2D are differentially expressed in thymic innate memory and virtual memory AIMT cells [2], a direct comparison of the gene expression between virtual memory and innate memory T cells has not been carried out.

Despite their abundance and functional uniqueness, AIMT cells are still understudied. The potential reasons for the underrepresentation of AIMT cells in the immunological research could be the confusing nomenclature [1] and unclear developmental and functional relationship between different types of AIMT cells. Moreover, only a minority of AIMT cell-related studies used germ-free mice [8, 16, 17], which might obscure the interpretation of the observations. It is unclear if the very existence of AIMT cells is physiological, because AIMT cells have been characterized exclusively in inbred mouse strains housed in clean conditions and because it has been proposed that the CD8^+^ T-cell compartment in laboratory mice substantially differs from feral mice [18].

Here, we show in side-by-side controlled experiments for the first time that the routes of AIMT cells formation are conserved in mice with different genetic backgrounds and hygienic conditions, including germ-free and feral mice.

## Methods

### Antibody

The antibodies used for flow cytometry are listed in Supplemental Table 1.

### Laboratory mice

Common laboratory mouse strains C57BL/6J and BALB/c and transgenic mouse strains Vβ5 [19] and C.Cg-Tg(DO11.10)10Dlo/J (henceforth DO11.10) [20] were used in this study.

Mice were bred in our facilities (SPF mice: Institute of Molecular Genetics of the Czech Academy of Sciences, Prague, Czech Republic and Department of Biomedicine, University Hospital Basel, Basel, Switzerland; germ-free mice: Laboratory of Gnotobiology, Institute of Microbiology of the Czech Academy of Sciences, Novy Hradek, Czech Republic) in accordance with laws of the Czech Republic and Switzerland, respectively. Animal protocols were approved by the Cantonal Veterinary Office of Baselstadt, Switzerland, by the Czech Academy of Sciences, the Czech Republic and by the respective Committees for the Protection and Use of Experimental Animals of the Institute of Microbiology (ID 5/2019) and of the Institute of Molecular Genetics (ID 8/2017, 15/2019) of the Czech Academy of Sciences.

Mice were kept in the animal facility with 12 hours of light and dark cycle with food and water ad libitum. Males and females were used for experiments. Age- and sex-matched pairs of animals were used in the experimental groups. If possible, littermates were equally divided into the experimental groups.

### Flow cytometry and cell counting

For the surface staining, cells were incubated with diluted antibodies in PBS/2% fetal calf serum on ice. LIVE/DEAD near-IR dye (Life Technologies, L34975) was used for discrimination of live and dead cells.

For the staining of feral mice, co-housed mice and control animals, cells were fixed with 2% formaldehyde at room temperature for 3 minutes prior to staining. All monoclonal antibodies used for this staining were tested for their compatibility with the fixation procedure.

For the intracellular staining, cells were fixed and permeabilized using Foxp3/Transcription Factor Staining Buffer Set (eBioscience, 00-5523-00). For some experiments, enrichment of CD8^+^ T cells was performed using magnetic bead separation kits EasySep (STEMCELL Technologies) or Dynabeads (Thermo Fisher Scientific) according to manufacturers’ instructions prior to the analysis or sorting by flow cytometry.

Cells were counted using Z2 Coulter Counter (Beckman) or using AccuCheck counting beads (Thermo Fisher Scientific) and a flow cytometer. Flow cytometry was carried out with a BD LSRII, BD Canto II or Cytek Aurora cytometers. Cell sorting was performed using Influx (BD Bioscience). Data were analyzed using FlowJo software (TreeStar).

### Stimulation of T cells

400,000 cells/ml freshly isolated lymph node cells in IMDM/10% fetal calf serum supplemented with IL-2 were stimulated with PMA (10 ng/ml) and ionomycin (1 μM), or IL-12 (10 ng/ml) and IL-18 (10 ng/ml), or left unstimulated for eight hours. 5 μg/ml Brefeldin A (Biolegend, 420601) was added to the culture for the last four hours of the incubation. The cells were stained with LIVE/DEAD near-IR dye and fluorescently labeled antibodies to CD8α, CD44, and CD49d. After washing, the cells were fixed and permeabilized and labeled with anti-IFN-γ antibody and analyzed by flow cytometry.

### Sublethal irradiation

C57BL/6J and BALB/c mice of the age 5-8 weeks were sublethally irradiated with 4 Gy on a T-200 X-ray machine (Wolf-Medizintechnik). The mice were sacrificed at day 16 post irradiation and the lymphocytes from the peripheral lymph nodes and the spleen were stained for CD122, CD8α, viability, CD44, CD62L, CD49d, CD4, TCRβ, Ly6c, and CD5 and analyzed by flow cytometry. The control group consisted of non-irradiated littermates.

### Adoptive transfer of cells into DO11.10 mice

Lymphocytes from peripheral lymph nodes and spleen from 8-12 weeks old BALB/c mice were harvested and stained with 5 μM Cell Trace Violet proliferation dye (Thermo Fisher, C34557) in PBS for 10 minutes at 37°C. 1×10^6^ naïve CD8^+^ CD44^-^ cells were sorted and i.v. injected into recipient DO11.10 mouse. The phenotype of the transferred cells was analyzed after 7 days.

### Samples for transcriptomic analysis

10-12 weeks old BALB/c or C57BL/6J SPF mice were irradiated with 4 Gy and sacrificed 16 days post irradiation. Non-irradiated mice were SPF young (7-12 weeks) or aged (12-19 months) BALB/c or C57BL/6J mice. Lymphocytes from the peripheral lymph nodes and the spleen were stained with TCRβ, CD8, CD44, CD49d, CD122 and viability dye and sorted by Influx. Each sample consisted of a pool of lymphocytes originating from 2-5 mice.

RNA was isolated using Trizol LS (Thermo Fisher Scientific) followed by in-column DNAse digestion using RNA clean&concentrator kit (Zymo Research) according to manufacturers’ instructions.

### Transcriptomic analysis

Samples were prepared with SMARTer^®^ Stranded Total RNA-Seq – Pico Input Mammalian library preparation kit v2 (Takara). Protocol included cDNA synthesis from mRNA, ligation of sequencing adaptors and indexes, ribosomal cDNA depletion, final PCR amplification and product purification. Library size distribution was evaluated on the Agilent 2100 Bioanalyzer using the High Sensitivity DNA Kit (Agilent). Libraries were sequenced using an Illumina NextSeq 500 instrument using 76 bp single-end high-output configuration.

Read quality was assessed by FastQC (http://www.bioinformatics.babraham.ac.uk/projects/fastqc). For subsequent read processing, a bioinformatics pipeline nf-core/rnaseq version 1.1 (https://doi.org/10.5281/zenodo.1447331), was used. Individual steps included removing sequencing adaptors with Trim Galore! (http://www.bioinformatics.babraham.ac.uk/projects/trim_galore/), mapping to reference genomes GRCm38 or Balb_cJ_v1 [21] (Ensembl annotation version 94) respective to the specimen strain with HISAT2 [22] and quantifying expression on gene level with featureCounts [23]. Resulting per gene read counts for individual strains were merged together using gene orthologue map obtained from Ensembl. Merged counts served as input for differential expression analysis using DESeq2 R Bioconductor package [24]. Prior to the analysis, genes not expressed in at least two samples were discarded. Shrunken log2-fold changes using the adaptive shrinkage estimator [25] were used for differential expression analysis. Genes exhibiting minimal absolute log2-fold change value of 1 and statistical significance of adjusted p-value < 0.05 between compared groups of samples were considered as differentially expressed for subsequent interpretation and visualization.

To show sample expression relatedness with the generic effect of mouse strain erased, we employed method of removing unwanted source of variation by fitting linear model using limma R Bioconductor package [26] (function removeBatchEffect) after normalizing the data using variance stabilizing transformation.

Raw fastq files have been deposited in the ArrayExpress database at EMBL-EBI (www.ebi.ac.uk/arrayexpress) under accession number E-MTAB-9091.

### Preparation of samples for TCR libraries

Lymphocytes from Vβ5 mice were sorted as viable CD8^+^ CD4^-^ TCRVα2^+^ CD62L^+^ CD44^-^ (naïve) or CD44^+^ (AIMT). RNA was isolated using Trizol (Thermo Fisher Scientific) followed by in-column DNAse treatment using RNA Clean&Concentrator kit (Zymo Research). cDNA synthesis was performed using a TCR specific primer (TTGGCACCCGAGAATTCCACTGHHHHHACAHHHHHACAHHHHHCACATTGATTTGGGA GTC) by using Maxima RT (Thermo scientific) according to manufacturer’s instructions. First round of PCR amplification was performed using primers CGTTCAGAGTTCTACAGTCCGACGATCHHHHACHHHHACHHHNGCAGCCCATGGTACCA GCAGTTCC and ACTGGAGTTCCTTGGCACCCGAGAATTC using HotStart Kappa HIFI ready mix (Kapa Biosystems) and the following conditions: 95°C for (5 min); 40 cycles: 98°C (20 s), 61°C (15 s), 72°C (15 s), and 72°C for (5 min). The resulting ~440 bp long PCR product was subjected to electrophoresis using an agarose gel and extracted using Gel Extraction Kit (Qiagen).

Following the first round of PCR, the TCRα copy number copies was quantified via qPCR (LightCycler 480 II, Roche) and compared to a dilution series of OT-I TCRα cloned in MSCV vector. 7.5×10^4^ copies of the PCR product were then used for the second PCR amplification round using primers AATGATACGGCGACCACCGAGATCTACACGTTCAGAGTTCTACAGTCCGACGAT*C and CAAGCAGAAGACGGCATACGAGATCGTGATGTGACTGGAGTTCCTTGGCACCC*G. The polymerase was HotStart Kappa HIFI ready mix (2x) and the following conditions: 95 °C (5 min); 40 cycles: 98 °C (20 s), 62 °C for (15 s), 72°C for (15 s), and 72 °C for (5 min). The resulting ~500 bp long PCR product was subjected to electrophoresis in agarose gels and purified using Gel Extraction Kit (Qiagen). The samples were mixed in equimolar ratios to obtain 30 μl of a final concentration of 10-nM and sequenced by Illumina MiSeq (v3 kit, 300bp paired-end reads) at the Biocev, Charles University, Vestec, Czech Republic.

### Analysis of TCR repertoires

FastQC was used to assess read quality. Next, MIGEC version 1.2.9 [27] was utilized for quantification of clonotypes and the resulting CDR3 sequence abundances in all samples were summarized to a clonotype abundance matrix. From this matrix, relative frequencies of CDR3 usage were computed. Clonotypes present in less than three samples were disregarded and the clonotype abundance matrix was treated similar to RNA-seq count matrix: DESeq2 was used to normalize for sequencing depth and RNA composition, and to test for differential clonotype abundance between AIMT and naïve sample groups. Normalized abundances were used for principal component analysis.

The sequence data has been deposited in the Sequence Read Archive and BioProject databases at NCBI (www.ncbi.nlm.nih.gov) under accession number PRJNA633838.

### Feral mice

We analyzed feral mice *Mus musculus musculus* that were trapped in human houses or garden shelters in the region of Central Bohemia, Czech Republic. The mice used in this study were feral mice caught up to one month prior to the analysis and 8-24 months old F1 offspring of the feral mice born in captivity. These mice were housed at the Faculty of Science, Charles University in Prague. Mice were provided with food *ad libitum* and were kept under stable conditions in open cages (13:11 hours, D:N, 23 °C).

All animal procedures on feral mice were carried out in accordance with the law of the Czech Republic paragraph 17 no. 246/1992 and the local ethics committee of the Faculty of Science, Charles University in Prague, specifically approved this study by accreditation no. 13060/2014-MZE-17214.

### Co-housing

Four SPF females were housed for eight weeks with one feral male per cage. The open co-housing cages were divided by a wire mesh with square openings (diameter was 1□cm), allowing contacts of co-housed animals. Co-housing cages were supplied with fresh bedding at the beginning of the experiment and provided with water and food *ad libitum*.

The age of SPF females was 4-6 weeks at the beginning of the co-housing experiment. The co-housed SPF females were matched with their littermates that remained in the SPF facility and had the same food and the same bedding as the co-housed animals. We performed two experiments with slightly modified setups. Experiment A: During the 6-week period of co-housing, the bedding was changed every 2 weeks and the feral males were transferred into another cage. Experiment B: During the 6-week period of co-housing, the bedding was not changed. Every 2 weeks, the mice were transferred to the other side of the same cage in order to be exposed to the feces of the feral mice.

### Collection of the samples for the microbiome analysis

A reference sample of saliva was collected from each mouse participating in the co-housing experiment before the start of the experiment B. Throughout both A and B experiments, the saliva has been collected from co-housed animals prior to the co-housing and after every second week of cohousing by gentle flushing with a pipette using 50□μ,l of the 0.9% NaCl solution. At the end of the experiment, the co-housed animals as well as the non-cohoused controls have been sacrificed and the samples from duodenum, jejunum, ileum, cecum and colon were collected for high-throughput 16S rRNA sequencing of the intestinal microbiota. At the end of each experiment, two co-housed Vβ5 animals and one co-housed feral male have been sent for a health status examination: HM FELASA Annually + additional agents (SOPF).

### Microbiota analysis by 16S rRNA sequencing

Metagenomic DNA from gut and saliva samples was extracted using the PowerSoil DNA kit (MO BIO Laboratories Inc., USA). Barcoding was performed with previously published 16S rRNA primers Bact-0341-b-S-17 and S-D-Bact-0785-a-A-21 [28]. Dual-indexed Nextera sequencing libraries were prepared using a previously published two step PCR design [29]. The first PCR was performed in 10 μl and consisted of 1x KAPA HiFi Hot Start Ready Mix (Roche), each 16S rRNA primer at 0.2 μM and 3-μl DNA. PCR conditions were as follows: initial denaturation at 95°C for 3 min followed by 28 (for colon and caecum), 33 (for saliva) or 35 (ileum, duodenum, jejunum) cycles each of 95°C (30 s), 55°C (30 s), and 72°C (30 s), and a final extension at 72°C (5 min). Second PCR differed from the first PCR in the following parameters: it was performed in 20 μl volume; the concentration of each indexed primer was 1 μM; 1μl (gut samples) or 6 μl (saliva) of the first PCR product were used as a template; and the number of PCR cycles was 12. Technical duplicates were prepared to account for noise due to PCR and sequencing stochasticity. PCR products were pooled equimolarly, size-selected with Pippin Prep (Sage Science, USA) at 520 - 750 bp and sequenced using MiSeq (Illumina, USA) and v3 chemistry (i.e. 2 × 300 bp paired-end reads). Sequencing data were archived in European Nucleotide Archive under project accession PRJEB38301. Accession numbers for each sample are presented in Supplemental Table 2.

Samples were demultiplexed and primers were trimmed by *skewer* software [30]. Using *dada2* [31], we filtered out low-quality sequences (expected number of errors per read > 1), denoised the quality-filtered fastq files and constructed abundance matrix representing reads counts for individual haplotypes (hereafter OTUs) in each sample. Using *uchime* [32] and gold.fna database we identified chimeric sequences and removed them from the abundance matrix. Taxonomic assignation of haplotypes was conducted by *RDP classifier* (80% confidence threshold, [33]) and Silva reference database (v 132, [34]).

Effect of cohousing on alpha diversity (based on Shannon diversity indices) was tested using linear mixed effect model (hereafter LMM), while considering individual identity as a random effect and the effect of treatment level (cohoused vs non-cohoused), gut section and their interactions as explanatory variables. The effect of cohousing on microbiota composition was assessed by permutation tests, where observed difference in composition (i.e. Bray-Curtis similarities) between cohoused lab vs. wild individuals and non-cohoused lab vs. wild individuals was compared with permuted average differences, where belonging to cohoused vs. non-cohoused lab group was randomized. OTUs, whose relative abundances were affected by cohousing were identified by *DESeq2* package [24].

## Results

### The frequency of AIMT cells increases in germ-free mice during aging

It has been shown that the percentage of AIMT cells increases upon aging [35–39]. However, these studies have used conventional specific pathogen free (SPF) mice and thus, the role of microbial antigens in this process cannot be excluded. Moreover, previous studies analyzed only C57BL/6 (henceforth B6) mice, but not BALB/c (henceforth Balb) mice that show substantial differences in the site of origin and cytokine dependency of AIMT cells [5, 40]. For this reason, we compared the CD8^+^ T-cell compartment in young and aged germ-free B6 and Balb mice. We observed that the frequency of peripheral CD44^+^ CD49d^-^ AIMT cells increases in both strains upon aging (Fig. 1A-B and S1A-B), suggesting a common homeostatic mechanism of age-dependent induction of AIMT cells.

**Figure 1.**
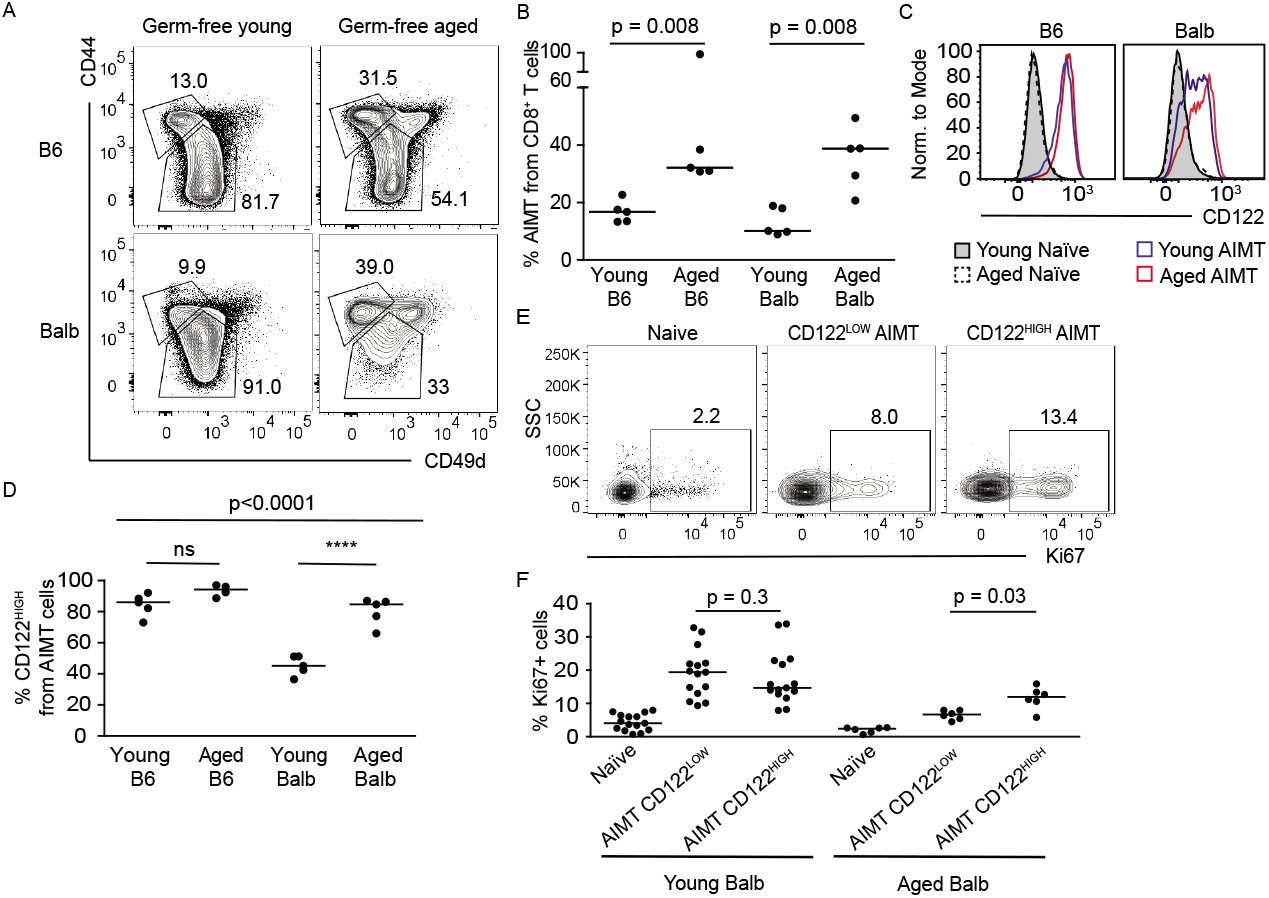
AIMT cells increase during aging in germ-free mice. (A-D) T cells from spleens of germ-free young (5-12 weeks) or aged (55-75 weeks) mice were stained for viability, TCRβ, CD8α, CD44, CD49d, and CD122 and analyzed by flow cytometry. (A) A representative staining of CD8^+^ T cells for CD44 and CD49d. Gates for naïve (CD44^-^) and AIMT (CD44^+^ CD49d^-^) cells are shown. A representative experiment out of 2 in total. (B) Quantification of the percentage of CD44^+^ CD49d^-^ AIMT cells among CD8^+^ T cells. n=5 mice from 2-5 independent experiments. Median is shown. The statistical significance was tested using 2-tailed Mann-Whitney test. (C) Histograms of CD122 expression in CD44^-^ naïve and CD44^+^ CD49d^-^ AIMT CD8^+^ cells from indicated mice. A representative experiment out of 2 in total. (D) Quantification of CD122^HIGH^ cells among CD8^+^ AIMT cells. Median is shown. The statistical significance was tested using ANOVA (p < 0.0001) with Bonferroni’s multiple comparison posttests of indicated pairs. **** p < 0.0001. (E-F) Splenocytes from young and aged SPF Balb mice were fixed and stained for CD8α, CD44, CD49d, CD122, and Ki67. (E) Ki67 expression in naïve, CD122^HIGH^ and CD122^LOW^ AIMT cells of aged mice. A representative mouse out of 6 in total. (F) Percentage of Ki67^+^ cells is shown. n= 6 (aged) or 15 (naïve) mice in 6 or 3 independent experiments, respectively. The statistical significance was tested using 2-tailed Wilcoxon matched-pairs signed rank test.

### Peripherally derived AIMT cells express high levels of CD122

AIMT cells from B6 depend on IL-15 and express high levels of CD122 (alias IL2RB), a subunit of the IL-15 receptor [5, 7, 16]. We observed that AIMT cells from young B6 mice express higher levels of CD122 than those from young Balb mice (Fig. 1C-D and S1C-D). Interestingly, a population of CD122^HIGH^ AIMT cells appears in aged Balb mice, whereas no difference in CD122 expression was found between AIMT cells from young and aged B6 mice (Fig. 1C-D). We observed that CD122^HIGH^ AIMT cells show slightly stronger homeostatic proliferation than CD122^LOW^ AIMT cells in aged, but not in young Balb mice (Fig. 1E-F), which probably contributes to their accumulation in aged mice.

CD122^HIGH^ AIMT cells in B6 are generated in the periphery via IL-15 and homeostatic TCR signaling [6, 17]. These cells are more frequent in aged mice, probably because the thymic output is low and the lymphopenic environment augments the AIMT cells formation. In contrast, CD122^LOW^ AIMT cells are generated in the thymus of young Balb mice via IL-4 [40]. We hypothesized that during aging of Balb mice, the thymic regress impairs de novo formation of CD122^LOW^ AIMT cells, leading to the lymphopenia-induced generation of proliferative CD122^HIGH^ AIMT cells in the periphery. To elucidate whether CD122 expression could discriminate between thymic (innate memory) and peripheral AIMT cells (virtual memory) in Balb mice, we employed two models for lymphopenia-induced AIMT cells formation.

The first lymphopenic model was sublethal irradiation of mice. Irradiation reduces the number of T cells, which leads to homeostatic proliferation and AIMT formation of the surviving T cells in the periphery [1]. As expected, irradiation increased the frequency of AIMT cells in B6 and Balb mice (Fig. 2A-B). Irradiation-induced AIMT cells showed significantly higher CD122 expression than control AIMT cells in Balb, but not in B6, mice (Fig. 2C-D). This result is in close agreement with the hypothesis that CD122 expression is a marker of peripherally induced AIMT cells in both strains.

**Figure 2.**
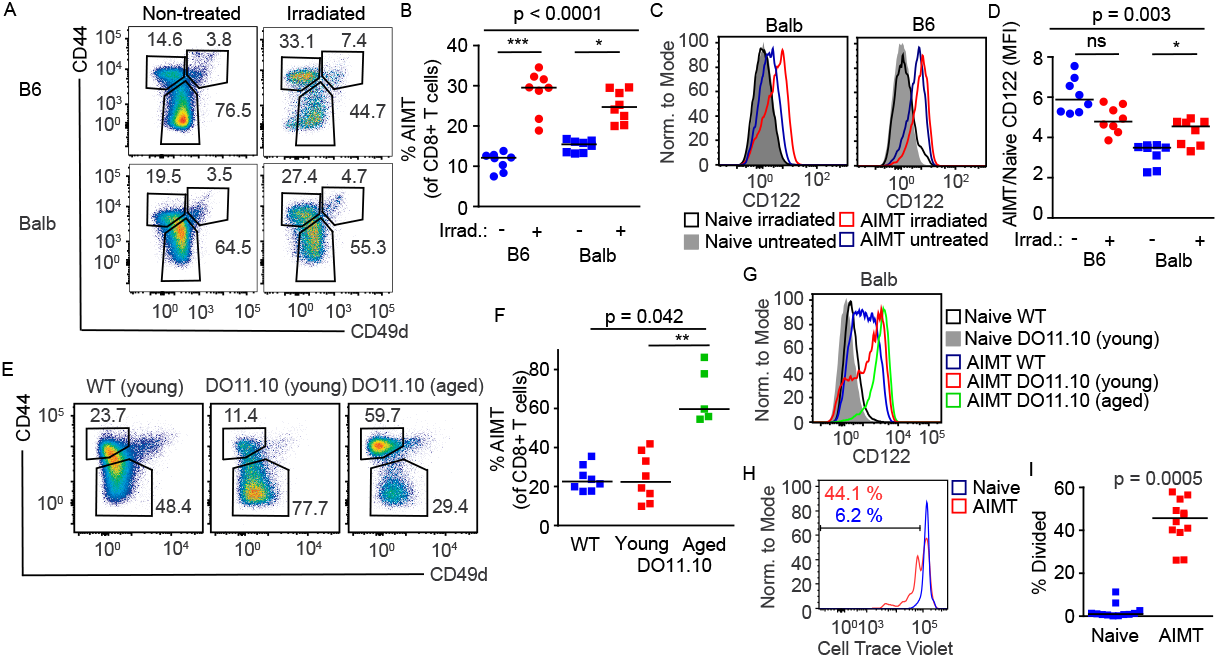
Lymphopenia induces CD122^HIGH^ AIMT cells in Balb mice. (A-D) 5-8 weeks old SPF B6 and Balb mice were sublethally irradiated or not. 16 days later, the mice were sacrificed and the splenocytes were stained for TCRβ, CD8α, CD44, CD49d, and CD122 and analyzed by flow cytometry. n=8 mice per group in 4 independent experiments. (A) A representative experiment out of 4 in total. (B) Quantification of the frequency of AIMT cells. The statistical significance was tested using Kruskall-Wallis test with Dunn’s multiple comparisons posttests. *** p < 0.001, * p <0.05. (C) Expression of CD122 in naïve and AIMT cells in untreated and irradiated B6 and Balb mice. (D) Quantification of CD122 expression in AIMT cells normalized to the expression on naïve T cells. (E-G) Splenocytes from young (5-14 weeks) Balb and young or aged (52-73 weeks) DO11.10 mice were stained for TCRβ, CD8α, CD44, CD49d, and CD122 and analyzed by flow cytometry. n=8 or 5 (aged DO11.10) mice in 4-5 independent experiments. (E) Representative experiments showing CD44 and CD49d staining of CD8^+^ T cells. (F) Quantification of the frequency of AIMT cells among CD8^+^ T cells. The statistical significance was tested using Kruskall-Wallis test with Dunn’s multiple comparisons posttests. ** p < 0.01 (G) Expression of CD122 in young Balb and young and aged DO11.10 naïve and AIMT CD8^+^ T cells. (H-I) Cell Trace Violet-loaded naïve CD8α^+^ CD44^-^ CD49d^-^ T cells isolated from lymph nodes and spleen from Balb mice were adoptively transferred into DO11.10 mice. 7 days later, the lymph nodes of the recipient mice were analyzed by flow cytometry. n=12 host mice in 3 independent experiments. (H) The Cell Trace Violet dye dilution in naïve (CD44^-^) and AIMT (CD44^+^ CD49d^-^) adoptively transferred cells (live TCRβ^+^ CD8^+^ TCRVβ8.1/8.2^-^) is shown. A representative experiment out of 3 in total. (I) The percentage of adoptively transferred naïve or AIMT cells that underwent at least one cycle of division is shown. The statistical significance was tested using Wilcoxon matched-pairs signed rank test.

The second model was TCR transgenic DO11.10 Rag^+^ Balb mice. Most of T cells in these mice are monoclonal CD4^+^ T cells specific to I-A^d^-OVA_323-339_ [41]. However, rare T cells expressing endogenous TCRα differentiate into CD8^+^ T cells. Thymic AIMT cells should be substantially reduced in these mice, because of the absence of IL-4-producing thymic iNKT cells that are crucial for the formation of thymic AIMT cells [40]. In addition, this mouse is lymphopenic in the CD8^+^ T-cell compartment, which should promote the formation of AIMT cells in the periphery.

DO11.10 mice showed comparable frequency of AIMT cells to polyclonal Balb mice. The frequency of AIMT CD8^+^ T cells during aging increased (Fig. 2E-F), indicating that age-dependent increase in the AIMT-cell compartment does not depend on the presence of iNKT cells. The expression of CD122 was higher in AIMT cells from DO11.10 mice than in AIMT cells from polyclonal Balb mice (Fig. 2G), which is consistent with the hypothesis that CD122^HIGH^ AIMT cells are formed in the periphery. When we transferred naïve polyclonal CD8^+^ T cells into the DO11.10 mice, we observed the formation of AIMT cells, which was coupled with their homeostatic proliferation (Fig. 2H-I).

Overall, these data indicate that CD122 expression differs between peripherally induced virtual memory AIMT cells (both in B6 and Balb mice) and thymus-derived innate memory AIMT cells of Balb mice.

### The site of origin and aging influence the gene expression profile of AIMT cells

The AIMT-cell compartment is heterogeneous. As mentioned above, AIMT cells are formed in different organs (thymus, peripheral lymphoid organs) and have distinct cytokine dependencies (IL-15 vs. IL-4) in different strains. Moreover, aging might change the gene expression programs of AIMT cells [36–38]. We addressed differences between particular AIMT-cell populations by high-throughput RNA sequencing using sorted naïve and AIMT cells from young B6 and Balb mice, irradiation-induced AIMT cells from B6 and Balb mice, AIMT cells from aged B6 mice, and CD122^LOW^ and CD122^HIGH^ AIMT cells from aged Balb mice.

As expected, the major source of the variability among these samples was the different genetic background of B6 and Balb mice (Fig. S2A). We normalized the gene expression data for the differences between the strains by fitting the data with a linear model using the strain identity as one of the explanatory variables (Fig. 3A). The expression of *Il2rb* (CD122) and *Itga4* (CD49d) in the sorted populations corresponded to the sorting strategy (Fig. 3B) and the previous characterization of AIMT cells from both mouse strains (Fig. 1A-D).

**Figure 3.**
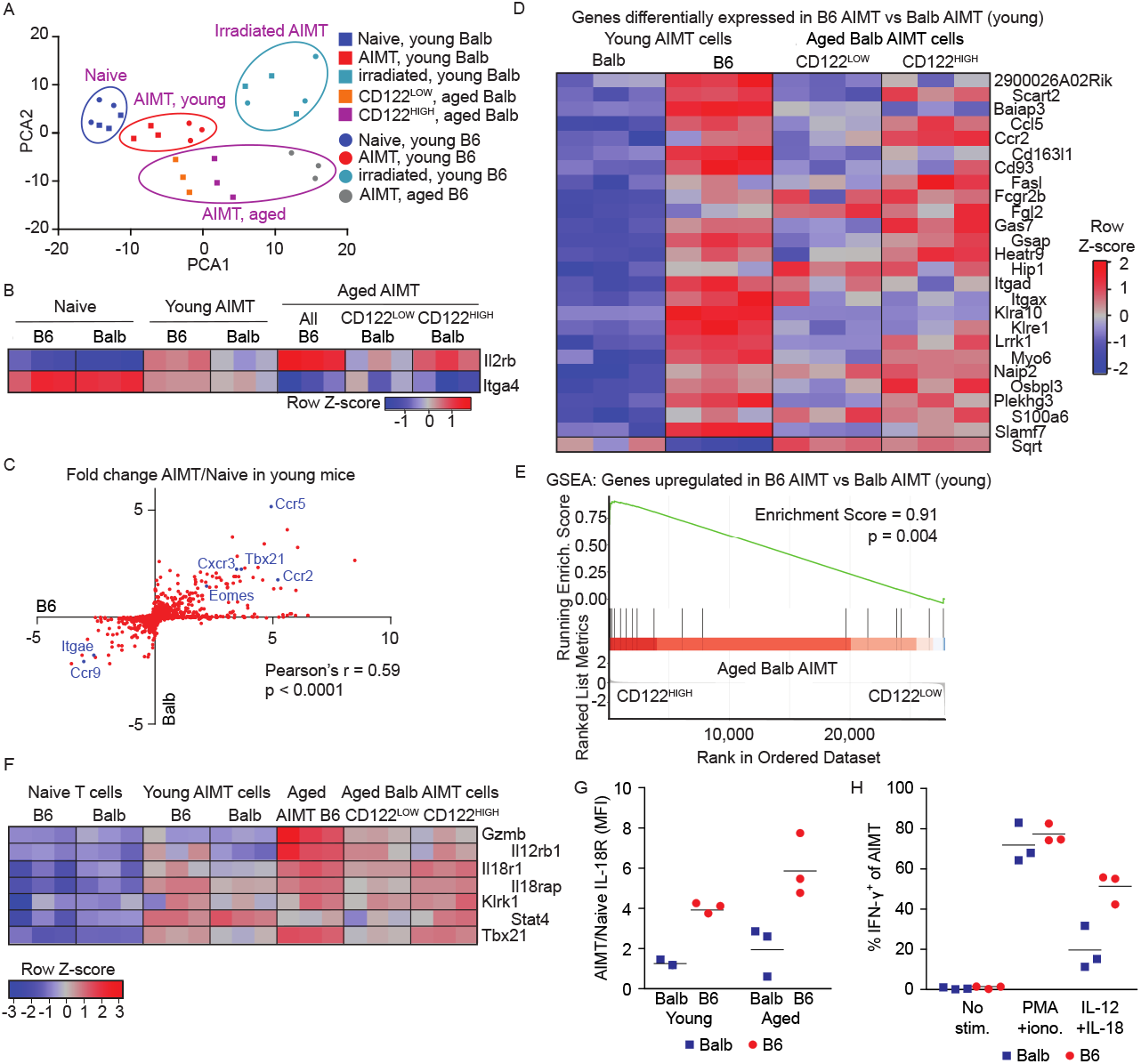
Similar gene expression profiles of thymic and peripheral AIMT cells. (A-E) Naïve (CD44^-^) and/or AIMT (CD44^+^ CD49d^-^) CD8^+^ T cells were sorted from lymph nodes and spleens from young (5-12 weeks) untreated or young irradiated (4 Gy, 3 weeks) B6 and Balb or aged B6 mice (76 weeks). CD122^HIGH^ and CD122^LOW^ AIMT cells were sorted from lymph nodes and spleens from aged Balb mice (54-55 weeks). The transcriptome of these populations was analyzed by RNA sequencing. (A) PCA analysis of the gene expression profiles of the individual samples normalized to avoid the strain-specific differences (top 500 variable genes). (B) Relative expression (z-score) of Il2rb (CD122) and Itga4 (CD49d) in individual samples using normalization for strain-specific differences. (C) Ratios of transcript levels in AIMT cells to naïve cells (young mice) were calculated for each gene and plotted as log2 for B6 and Balb strains. Selected genes are indicated. The correlation was calculated using Pearson correlation coefficient. (D) Relative expression (z-score) of genes showing significantly different expression between AIMT cells from young B6 and Balb mice (after the strain normalization) in AIMT cells from B6 and Balb mice and in CD122^HIGH^ and CD122^LOW^ AIMT cells from aged Balb mice. (E) The gene set enrichment analysis of genes with significantly higher expression in B6 than in Balb mice (D) in CD122^HIGH^ vs. CD122^LOW^ AIMT cells from aged Balb mice. (F) Relative expression (z-score) of selected genes involved in by-stander cytotoxicity between AIMT cells from young B6 and Balb mice (after the strain normalization) in AIMT cells from B6 and Balb mice and in CD122^HIGH^ and CD122^LOW^ AIMT cells from aged Balb mice. (G) Expression of IL-18R on the surface of young and aged Balb and B6 AIMT cells measured by flow cytometry. Geometric mean fluorescent intensity (MFI) was normalized to naïve cells in the same sample. (H) Production of IFN-γ by AIMT cells (gated as CD8α’ CD44^+^ CD49d^4^) from young adult B6 or Balb mice was measured by flow cytometry. Lymph node cells were stimulated with PMA + ionomycin or with IL-12 and IL-18 or left unstimulated for 8 hours and analyzed by flow cytometry.

We assumed that naïve and irradiation-induced AIMT cells represent two relatively uniform cell types present in both strains, and we used them as controls for evaluating our data normalization. The principal component analysis (PCA) of the adjusted data revealed that naïve cells and irradiation-induced AIMT cells from both strains clustered together, confirming the reliability of our adjusted data (Fig. 3A). We obtained very similar results when performing the PCA using data without strain-specific normalization, but with the exclusion of significantly differentially expressed genes between the strains (Fig. S2B),

The AIMT cells from aged mice showed moderate differences in their gene expression patterns in comparison to AIMT cells from young animals (Fig. 3A). We identified 36 genes upregulated in AIMT cells from aged mice vs. young mice in both strains (Fig. S2C). Twenty four of these transcripts were at least 4-fold enriched in AIMT cells from aged B6 mice in comparison to young mice in the previous study by Quinn et al. [36]. Interestingly, we did not identify a single gene significantly upregulated in AIMT cells from young mice vs. aged mice in both strains.

Young AIMT cells from both strains showed comparable global gene expression signature. The gene expression similarity between these populations was revealed as the proximity of AIMT young cells from both strains in the PCA plots (Fig. 3A, Fig. S2B) as well as a correlation of AIMT/naïve cell fold changes for individual genes plotted as B6 vs. Balb without using any correction for the strain-dependent differences (Fig. 3C). Despite the similarity of the gene expression patterns, we could identify one gene significantly upregulated, and 25 genes significantly downregulated in young Balb AIMT cells vs. B6 AIMT cells (Fig. 3D), suggesting that these genes might distinguish thymic innate memory and peripheral virtual memory subsets of AIMT cells.

We observed that the genes upregulated in AIMT cells from young B6 vs. young Balb mice are significantly enriched in aged CD122^HIGH^ AIMT cells vs. aged CD122^LOW^ AIMT cells from aged Balb mice (Fig. 3D-E), supporting the hypothesis that high CD122 levels predict the peripheral origin of AIMT cells irrespective of the strain.

It has been proposed that T-bet (*Tbx21*) and NKG2D (*Klrk1*) are expressed in peripheral, but not in thymic AIMT cells [2]. Moreover, a small group of genes potentially involved in by-stander cytotoxicity (*Gzmb, Il12rb2, Il18r1, Il18rap, Stat4*) were shown to be upregulated in peripheral AIMT cells [2, 6]. All of these genes, but *Gzmb*, were enriched in young AIMT cells over naïve T cells in our data (Fig. 3F). Four genes (*Tbet, Klrk1, Il18r1, Il18rap*) showed higher expression in AIMT cells of the proposed peripheral origin (AIMT cells from young and aged B6, and CD122^HIGH^ AIMT cells from aged Balb mice) than in the corresponding AIMT cells of the proposed thymic origin (AIMT cells from young Balb and CD122^LOW^ AIMT cells from aged Balb mice) (Fig. 3F).

Flow cytometry confirmed higher expression of IL-18R in AIMT cells from B6 than from Balb mice (Fig. 3G, Fig. S2D). Moreover, IL-18R was preferentially expressed in CD122^high^ AIMT cells (Fig. S2E). Accordingly, AIMT cells from B6 mice were more efficient in producing IFN-γ after stimulation with IL-12/IL-18 than AIMT cells from Balb mice, whereas the responses of these cells to PMA and ionomycine were comparable (Fig. 3H, Fig. S2F).

These data are consistent with our hypothesis that CD122 expression can discriminate between thymic and peripheral T cells. Moreover, these data indicate that thymic innate memory and peripheral virtual memory AIMT cells differ in their ability to facilitate by-stander immune protection by responding to pro-inflammatory cytokines.

### Young and aged AIMT cells represent identical TCR clonotypes

The frequency of murine AIMT cells changes in time. It peaks around 3 weeks of age [42] and it increases again as the mouse ages [35]. For this reason, it is unclear whether AIMT cells in young and aged animals have a different clonal origin [43]. To address this question, we analyzed TCR sequences of naïve and AIMT cells in young and aged mice. To overcome the problem with TCRα and TCRβ pairing, we analyzed TCRα chains in Vβ5 mouse that has a fixed TCRβ from the OT-I TCR. The frequency of AIMT cells in Vβ5 mice recapitulates WT mice, including the increase in the frequency of AIMT cells upon aging (Fig. S3A).

First, we quantified the usage of TRAV14 (Vα2) and TRAV12 (Vα8) by staining with specific antibodies. In line with our previous data [17], we observed a higher Vα2/Vα8 ratio in AIMT cells than in naïve T cells of young mice (Fig. 4A, S3B). Interestingly, the Vα2/Vα8 ratio in naïve cells was higher in aged mice than in young mice (Fig.S3B). The highest Vα2/Vα8 ratio was observed in the AIMT compartment in aged mice. Because Vα2^+^ T cells are on average more self-reactive than Vα8^+^ T cells in Vβ5 mice [17], our data suggest that the CD8^+^ T-cell compartment in aged mice is enriched for self-reactive T cells and that AIMT cells are generally more self-reactive than naïve cells in young as well as in aged mice.

**Figure 4.**
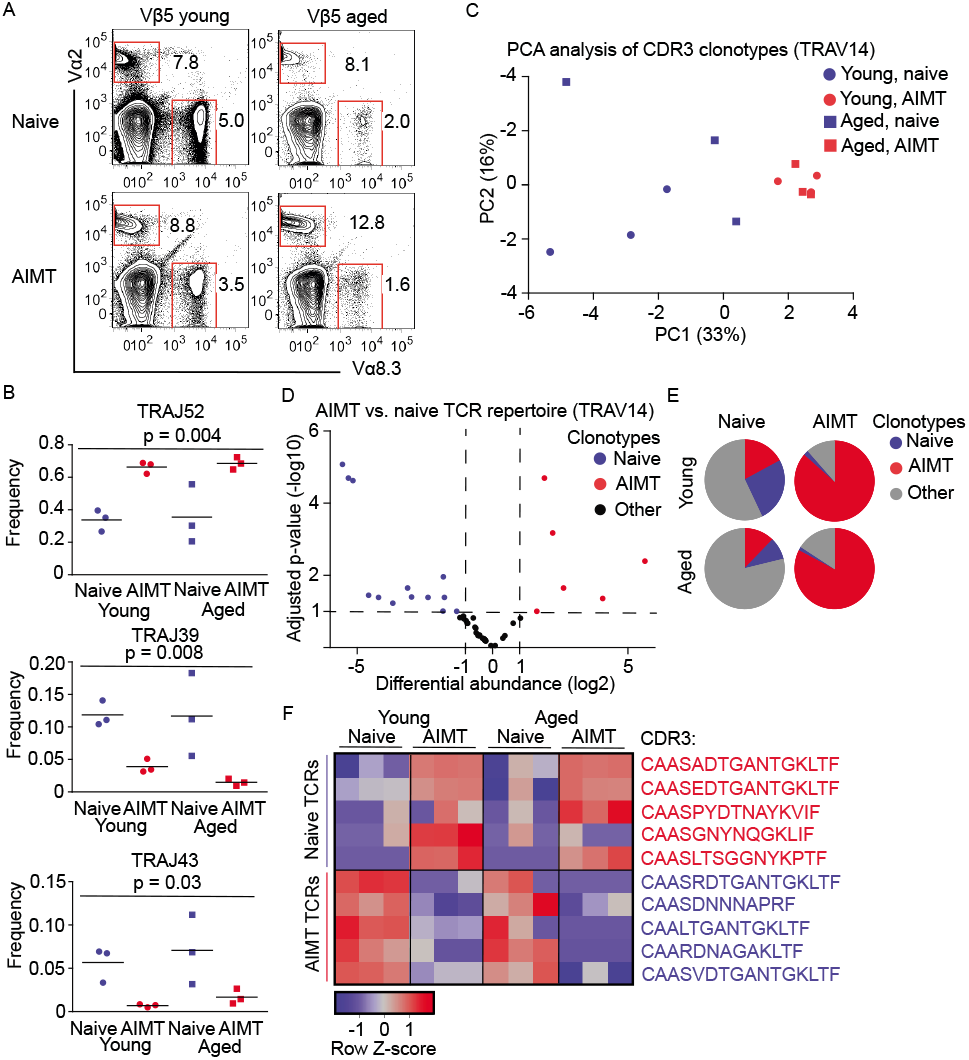
AIMT cells in young and aged mice represent identical clones. (A-F) Lymph node cells from young (7-12 weeks) and aged (65-77 weeks) Vβ5 mice were enriched for CD8^+^ T cells by magnetic beads and stained for CD8, CD44, CD62L, CD49d, TCRVα2 and TCRVα8.3. 2-3 mice were pooled for each sample. 3 independent experiments. (A) The percentage of TCRVα2’ and TCRVα8.3’ among AIMT and naïve CD8^+^ T cells is shown. (B-F) AIMT (CD44’ CD62L^+^) and naïve (CD44^-^ CD62L’) TCRVα2’ CD8^+^ T cells were FACS sorted. Library of TCR sequences was generated from isolated RNA using a two-step PCR and analyzed by deep sequencing. (B) Frequency of TRAJ52, TRAJ39, and TRAJ43 usage among the analyzed samples. The statistical significance was tested using one-way repeated measures ANOVA. (C) PCA analysis of the TCR clonotype (based on CDR3 amino acid sequence) usage in the individual samples. Only clonotypes detected in at least 3 samples were included in the analysis. (D) A volcano plot showing log2 differential usage between naïve and AIMT cells (both young and aged) and adjusted p-value for each individual TCR clonotype present at least in 3 samples. “AIMT”, “naïve” (2-fold difference in usage, adjusted p<0.01) and “neutral” (non-significant) clonotypes are indicated in red, blue, and gray respectively. (E) Pie charts showing the abundance of the “AIMT”, “naïve”, and neutral clonotypes in the T-cell repertoires of young and aged Vβ5 mice. (F) Relative usage (z-score) of 5 most abundant “AIMT” and 5 most abundant “naïve” clonotypes in the indicated samples.

In the next step, we analyzed TCRs from Vα2^+^ T cells in greater detail using high-throughput sequencing. Naïve and AIMT cells showed differential usage of TRAJ segments, indicating that they are formed from different T-cell clonotypes (Fig. 4B, S3C), which is in line with previous results [8, 17]. Interestingly, we did not see any differences in the TRAJ usage between young and aged mice (Fig. 4B, S3C). The second step of our analysis was the comparison of individual clonotypes using CDR3 amino acid sequences. The PCA analysis separated a cluster of AIMT cells (both young and aged mice) from naïve T cells (Fig. 4C). The comparison of the usage of individual clonotypes by naïve and AIMT cells showed that some clonotypes are enriched in the naïve compartment, whereas some clonotypes are significantly enriched in AIMT cells (Fig. 4D). The AIMT cell repertoire in young and aged mice is constituted predominantly from a small number of AIMT cell-enriched clonotypes. In contrast, the naïve repertoire is more diverse and contains naïve-enriched as well as AIMT-enriched clones (Fig. 4E). The usage of the dominant clonotypes differ in naïve and AIMT subsets, but not so much between the corresponding subsets from young and aged mice (Fig. 4F). Overall, these data showed that AIMT cells from young and aged mice have the same clonal origin.

### AIMT cells in feral mice

It has been suggested that the exposure to environmental antigens in feral and ‘pet shop’ mice has a substantial role in shaping the CD8^+^ T-cell compartment [18]. To address the possibility that the occurrence of AIMT cells might be an artifact of specific hygienic conditions in germ-free and SPF facilities, we bred feral mice in the animal facility and analyzed F1 offspring born in captivity. Interestingly, we observed that AIMT cells are relatively frequent in these mice (Fig. 5A, S4A). The expression of CD122 was variable between individual animals, suggesting variable peripheral or thymic origin of these cells likely based on the genotype of the mice.

**Figure 5.**
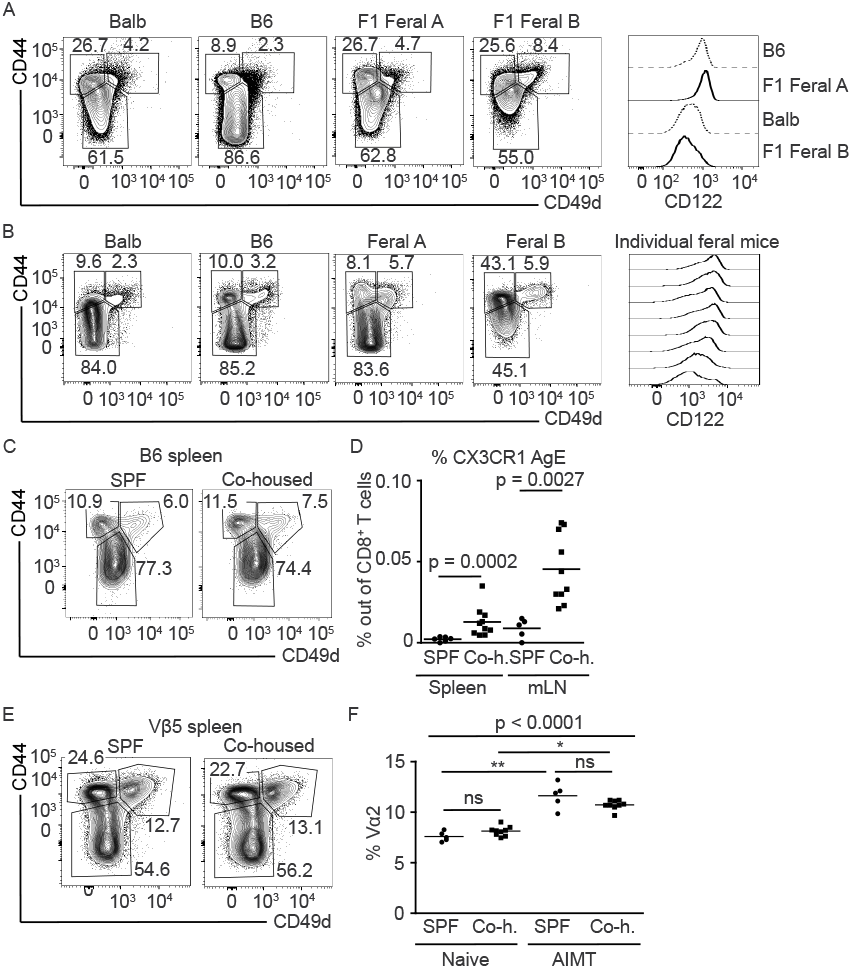
AIMT cells are present in feral mice and in laboratory mice co-housed with feral mice. (A-B) Spleens of laboratory B6 and Balb strains and 8-24 months old F1 offspring of feral mice (A) or feral mice (B) were isolated, fixed with formaldehyde and stained for CD8, CD44, CD49d, and CD122 and analyzed by flow cytometry. Percentage of naïve (CD44^-^), AIMT (CD44^+^ CD49d^-^), and antigen experienced (CD44^+^ CD49d^+^) cells among CD8^+^ T cells and CD122 expression in AIMT cells are shown. (C-F) B6 and Vβ5 mice were co-housed with feral mice or F1 offspring of feral mice for 6 weeks and their splenocytes were analyzed by flow cytometry. (C) Percentage of naïve (CD44^-^), AIMT (CD44^+^ CD49d^-^), and antigen experienced (CD44^+^ CD49d^+^) cells among CD8^+^ T cells in SPF and co-housed B6 mice. A representative experiment is shown. (D) Percentage of CX3CR1^+^ antigen experienced cells among all CD8+ T cells is shown. Statistical significance was calculated using Mann-Whitney test. (C-D) n = 5 (SPF) or 10 (co-housed) mice in 2 independent experiments. (E) Percentage of naïve (CD44^-^), AIMT (CD44^+^ CD49d^-^), and antigen experienced (CD44^+^ CD49d^+^) cells among CD8^+^ T cells in SPF and co-housed Vβ5 mice. A representative experiment is shown. (F) Percentage of TCRVα2^+^ T cells among AIMT (CD44^+^ CD49d^-^) and naïve (CD44^-^) CD8^+^ T cells from SPF and co-housed Vβ5 mice. Statistical significance was tested using Kruskal-Wallis test with Dunn’s multiple comparison posttests.

In the next step, we analyzed feral mice trapped in garden shelters and human houses. We observed that these mice contain a large population of CD8^+^ CD44^+^ CD49d^-^ T cells, corresponding to AIMT cells (Fig. 5B, S4A). The expression of CD122 in these AIMT cells varied among individual animals, indicating the role of genetic factors (Fig. 5B). Overall, these data suggested that the population of AIMT cells exists in feral mice.

### Co-housing of laboratory and feral mice has minimal effects on the CD8^+^ T-cell compartment

To observe how the AIMT-cell compartment and other CD8^+^ T-cell subsets change upon exposure to environmental antigens in laboratory mice, we co-housed B6 WT mice and transgenic Vβ5 mice with feral mice for 6 weeks. Surprisingly, the co-housing had no significant effect on the proportions of the major CD8^+^ T-cell subsets, i.e., naïve, AIMT, and antigen-experienced cells, in B6 and Vβ5 mice (Fig. 5C-E, S4B-D). The only observed difference between co-housed and control laboratory strains was a slightly higher percentage of CX3CR1^+^ CD44^+^ CD49d^+^ antigen-experienced cells in the spleen and mesenteric lymph nodes of the co-housed animals of the SPF origin (Fig. 5D, S4C).

Co-housing with feral mice did not significantly alter the composition of the CD8^+^ compartment of Vβ5 mice (Fig. S4D). We observed that the enrichment of the AIMT-cell compartment in Vα2^+^ T cells was preserved in the co-housed animals (Fig. 5F, S4E), suggesting that the co-housing did not substantially alter the composition of the repertoire of the AIMT-cell compartment. Overall, our data indicate that exposure to environmental antigens has a minimal impact on the AIMT-cell compartment.

### Limited effects of co-housing on the microbial colonization

Because we observed a surprisingly small effect of the co-housing of laboratory strains with feral mice on their CD8^+^ T-cell compartment, we examined the colonization of the feral and co-housed animals with microbes and viruses. A detailed health examination of a limited number of animals detected *Helicobacter, Pasteurella*, MCMV, and endoparasites, but no other pathogenic agents, in the feral mice (Appendix 1). Of these pathogens, only *Helicobacter* and endoparasites were transmitted to the co-housed animals in at least 1 case out of 4 tested co-housed mice (Appendix 1). This was in line with the fact that none of the co-housed mice died before the end of the experiment, including Vβ5 mice, which are most likely immunocompromised because of the limited TCR repertoire.

Subsequently, we addressed the effects of co-housing on bacterial colonization. We performed 16S rRNA sequence analysis of the composition of the microbial community in the gut and salivary glands of co-housed animals. The overall microbial diversity in the intestine of co-housed laboratory strains was not affected except for the ileum, where the diversity increased (Fig. S5A). Interestingly, microbial diversity decreased in the colon of feral mice upon co-housing (Fig. S5A). As expected, the microbial composition of the intestine of the laboratory strains converged to the feral mice upon cohousing (Fig. 6A, Fig. S5B). The most notable effect of the co-housing was the abundance of Campylobacteria in the colon and caecum of the co-housed mice, whereas this group of bacteria was not present in non-co-housed laboratory mice (Fig. 6B, Fig. S5C). Two different *Helicobacteria* taxons (Campylobacterales) and *Alistipes* (Bacteroidia) were detected in the colon and caecum of some of the co-housed, but not SPF, mice (Fig. 6C). In contrast, three taxonomic units from Clostridiales, *Roseburia*, and *Ruminococcus* taxons were depleted in colon, caecum and duodenum of the co-housed mice (Fig. S5D).

**Figure 6.**
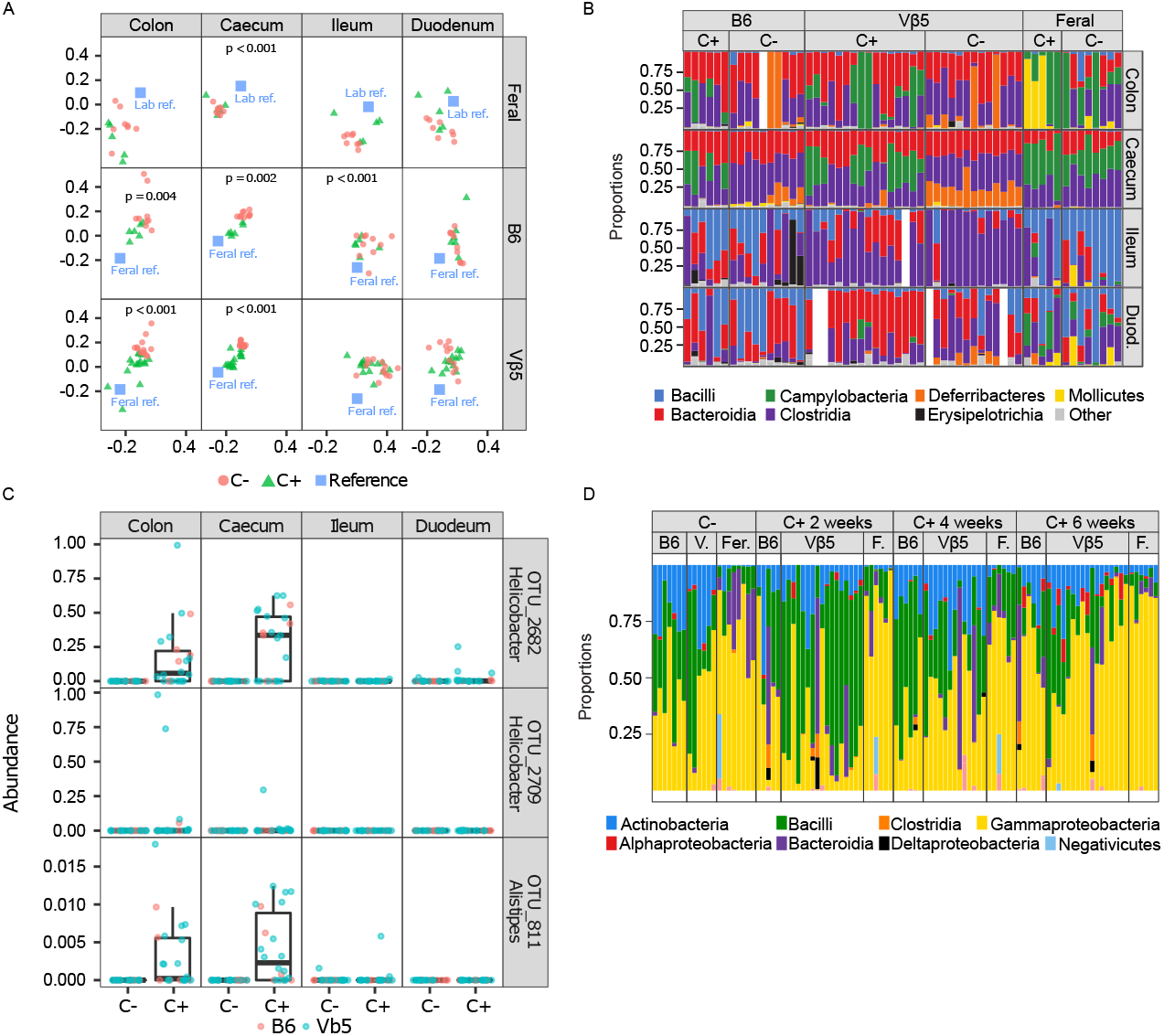
Co-housing with feral mice changes the intestinal microbial colonization of laboratory mice. (A-C) The analysis of intestinal microbiome of co-housed and non-co-housed laboratory and feral mice (Experiment B, see Methods). Samples were taken from the colon, caecum, ileum, or duodenum. C+; co-housed mice for 6 weeks (laboratory + feral), C-; non-co-housed mice. Duod. – Duodenum. (A) Compositional variation in cohoused vs. non-cohoused groups assessed based on Nonmetric Multidimensional Scaling (NMDS) of Bray-Curtis dissimilarities. Centroids for a reference group (feral non-cohoused mice in the case B6 or Vβ5 and non-cohoused Vβ5 plus B6 in the case of feral mice) are indicated by blue rectangles. Permutation-based p-values indicate significant changes due to cohousing. (B) Taxonomical composition of the intestinal microbiota of laboratory co-housed and non-co-housed B6, Vβ5, and feral mice. Color bars represent proportions of dominant bacterial classes in each sample. White space means no data. (C) Three most abundant operational taxonomic units with higher relative abundance in co-housed than non-co-housed B6 and Vβ5 laboratory mice. (D) Taxonomical composition of the salivary microbiota of laboratory B6 or Vβ5 mice co-housed (C+) vs. non-co-housed (C-) with feral mice (Experiment B, see Methods). Color bars represent proportions of dominant bacterial classes in each sample. Samples were collected from each mouse prior to the co-housing (C-) and after 2, 4, and 6 weeks during the co-housing experiment. F – feral.

The salivary microbiome of the laboratory strains did not converge to feral mice, but rather the microbiome of the feral mice converged to the microbiome of SPF laboratory strains during the cohousing (Fig. S5E). However, co-housing led to a dramatic increase in Bacteriodia taxon in saliva of laboratory strains (Fig. 6D, Fig. S5F). Moreover, the abundance of Clostridia and Deltaproteobacteria was detected in some individuals during, but not before, co-housing (Fig. 6D, Fig. S5F). Overall, these data document that co-housing with feral mice changed the microbial colonization of laboratory mice without a substantial impact on the CD8^+^ T-cell compartment.

## Discussion

AIMT cells represent one of the two most numerous populations of CD8^+^ T cells in mice. Although it has been shown that these cells are functionally different from naïve and antigen-experienced memory T cells [6, 17, 39, 44–47], our understanding of the biology of these cells is incomplete. There are multiple potential reasons why AIMT cells are an underrepresented area of focus. First, multiple ways of AIMT cells formation were proposed involving different cytokines (IL-4, IL-7, and IL-15), different anatomical sites (thymus, secondary lymphoid organs), and different physiological states (experimentally induced lymphopenia, newborn lymphopenia, aging) [1, 2, 4]. Second, the phenotypic and functional similarities between particular AIMT cell subsets and their differences are not well understood [1, 2, 4]. Third, only a minority of studies used germ-free mice to exclude microbial antigens [16, 17] and at the same time, no previous study addressed the presence of AIMT cells in feral mice, which were reported to have substantially different CD8^+^ T cell compartment from laboratory mice [18].

In this study, we performed comprehensive phenotyping of AIMT cells to address multiple unresolved questions concerning their origin and heterogeneity. We concluded that the formation of AIMT cells represent a robust genetic differentiation program across the genetic background, aging, and hygienic status. However, these three conditions regulate AIMT cell abundance and internal variability.

We identified CD122 as a marker discriminating CD122^LOW^ thymic (innate memory) and peripheral CD122^HIGH^ (virtual memory) subsets of CD8^+^ AIMT cells. CD122^LOW^ AIMT cells prevail in WT young Balb mice and some feral mice. On the contrary, CD122^HIGH^ AIMT cells are dominant in B6, in some feral mice and Balb mice with the impaired formation of CD122^LOW^ cells because of aging, expression of a transgenic TCR disabling the formation of IL-4 producing iNKT cells, or irradiation. Collectively, our data indicate that the absence of thymic innate memory AIMT cells allows the conversion of naïve T cells into AIMT cells in the periphery, presumably via IL-4, IL-7, and/or IL-15 [1, 4, 7, 11, 12].

AIMT cells represent the largest population of CD8^+^ T cells in aged B6 mice [35]. Here we confirmed this observation in germ-free conditions and extended it by the analysis of aged Balb mice. The mechanism behind the high frequency of AIMT cells in aged animals was unclear. It has been proposed that the origin of these cells might differ from the AIMT cells in young animals [37]. We addressed this question by analyzing gene expression profiles and TCR profiling of young and aged AIMT cells using high-throughput sequencing. In addition to confirming differences between AIMT cells from young and aged B6 mice as reported previously [36], we identified similarities between CD122^HIGH^ AIMT cells from aged Balb mice and AIMT cells from B6 mice. These similarities support the concept that CD122^HIGH^ AIMT cells are formed at the periphery in both B6 and Balb strains. Expression of key effector molecules involved in by-stander cytotoxicity as well as IL-12/IL-18 production of IFN-γ was higher in peripheral AIMT cells than in thymic AIMT cells, suggesting that these two subsets differ in their ability to elicit by-stander protective immunity as proposed previously [2].

In line with recent studies [8, 17], we identified that CD8^+^ naïve and AIMT cells use different TCR repertoires. We observed that clones typically present in the naïve population are virtually absent in AIMT cells, whereas typical AIMT clones can still be identified in the naïve compartment, albeit with a much lower frequency. This partial overlap is not surprising as AIMT cells are formed from naïve T cells in the periphery. However, a recent study by Miller et al. showed a clear separation of naïve and AIMT-cell repertoires [8]. This discrepancy might be caused by the difference between the employed mouse models. We analyzed TCRVα2 repertoire in transgenic mice expressing TCRVβ5 adopted from H2-K^b^-restricted OT-I TCR. Importantly, the Vβ5 mouse has a similar frequency of AIMT cells as WT B6 mice. Miller et al. analyzed TCRα repertoire from mice expressing TCRVβ3 chain. This chain usage is abundant among prostate cancer-infiltrating regulatory CD4^+^ T cells [48] and might produce T-cell repertoire biased towards MHCII-restriction, subsequently affecting the frequency, repertoire or phenotype of CD8^+^ AIMT cells.

We observed that young and aged AIMT cells have identical TCRVα2 repertoire, implying that the AIMT compartment in young and aged mice consists of the same T-cell clones. Moreover, we observed that naïve and AIMT cells in aged mice preferentially use TCRVα2 over TCRVα8.3 chains, which most likely reflects the survival and proliferation advantage of relatively highly self-reactive TCRVα2^+^ T-cell clones during aging, which is in good agreement with a previous report [49].

Overall, the probable causes of the high frequency of AIMT cells in aged B6 and Balb mice are: (i) low thymic egress of fresh naïve T cells, (ii) irreversible AIMT cells formation from highly self-reactive clones augmented by age-related lymphopenia, and (iii) the survival and proliferative advantage of AIMT cells over naïve T cells.

For the first time, we studied AIMT cells in feral mice and F1 offspring of feral mice born in captivity, suggesting that AIMT cells are not an artifact of inbred strains and/or SPF conditions. Feral mice showed an even higher frequency of AIMT cells than laboratory strains. This could be caused by the enhancement of the AIMT-cell compartment via IL-4 produced in response to helminth [46, 47] or other infections. Although it has been shown that co-housing of laboratory strains with ‘pet shop’ mice leads to dramatic changes of their CD8^+^ T cell compartment [18], we could detect only a minor increase of CX3CR1^+^ effector cells in co-housed B6 mice. Otherwise, the CD8^+^ T-cell compartment in the spleen, peripheral lymph nodes, and mesenteric lymph nodes of co-housed mice was unaffected. A possible explanation is that our feral mice showed much lower pathogenic burden than the ‘pet shop’ mice used by Beura et al. [18], which seems logical given the high animal density and the lack of natural predation at ‘pet shop’ farms. Although different populations of feral mice might have a different pathogenic burden, our data document that co-housing with feral mice does not always induce dramatic changes in the CD8^+^ T cell compartment. We observed that co-housing with feral mice substantially changes the microbial composition of the intestine and saliva of laboratory strains, including opportunistic pathogens such as *Pasteurella* and *Helicobacter*.

Altogether, our data document that AIMT cells represent a large CD8^+^ T-cell compartment in mice, which is present in germ-free as well as in feral conditions. The frequency of peripherally induced self-reactive CD122^HIGH^ AIMT cells increases during aging via homeostatic mechanisms in Balb and B6 mice.

## Acknowledgement

We thank Ladislav Cupak for technical assistance and genotyping of mice and Sarka Kocourkova for preparation of RNAseq libraries. This study was supported by the Czech Science Foundation (19-03435Y to OS, 18-17796Y to JKr, 19-02261S to HK), Swiss National Science Foundation (Promys, IZ11Z0_166538 to OS), and the European Research Council (FunDiT to OS). The Laboratory of Adaptive Immunity was supported by an EMBO Installation Grant (3259 to OS). The Laboratory of Adaptive Immunity and the Laboratory of Genomics and Bioinformatics were supported by the Institute of Molecular Genetics of the Czech Academy of Sciences core funding (RVO 68378050). OS was supported by the Purkyne Fellowship provided by the Czech Academy of Sciences. PS, RS, and TM were supported by MICOBION project funded from EU H2020 (No 810224). JN and MK were supported by “Center for Tumor Ecology - Research of the Cancer Microenvironment Supporting Cancer Growth and Spread” OP RDI CZ.02.1.01/0.0/0.0/16_019/0000785 provided by the Czech Ministry of Education, Youth and Sports and the European Regional Development Fund. LS was supported by the Charles University specific research grant (SVV 260571/2020). AM and VN were students partially supported by the Faculty of Science, Charles University, Prague. JN was a student partially supported by the University of Chemistry and Technology, Prague.

The animal facility of the IMG is a part of the Czech Centre for Phenogenomics and the work there was supported in part by following grants: LM2015040, OP RDI CZ.1.05/2.1.00/19.0395, OP RDI BIOCEV CZ.1.05/1.1.00/02.0109 provided by the Czech Ministry of Education, Youth and Sports and the European Regional Development Fund.

## Author contribution

AM performed most of the experiments. AS, LS, MP, AD, RS, TM, AN, KK, PS, OS performed experiments. AM, VN, and OS analyzed data and finalized figures. JKu and JM analyzed the transcriptomic data. JN analyzed the TCR profiling data. DC and JKr analyzed the S16 sequencing data. RS and PS provided feral mice. TH and HK provided germ-free mice. MK, HK, JKr, PS, and OS supervised the work. OS conceived the study. AM, VN, and OS wrote the manuscript. All the authors contributed to the manuscript and reviewed it.

---

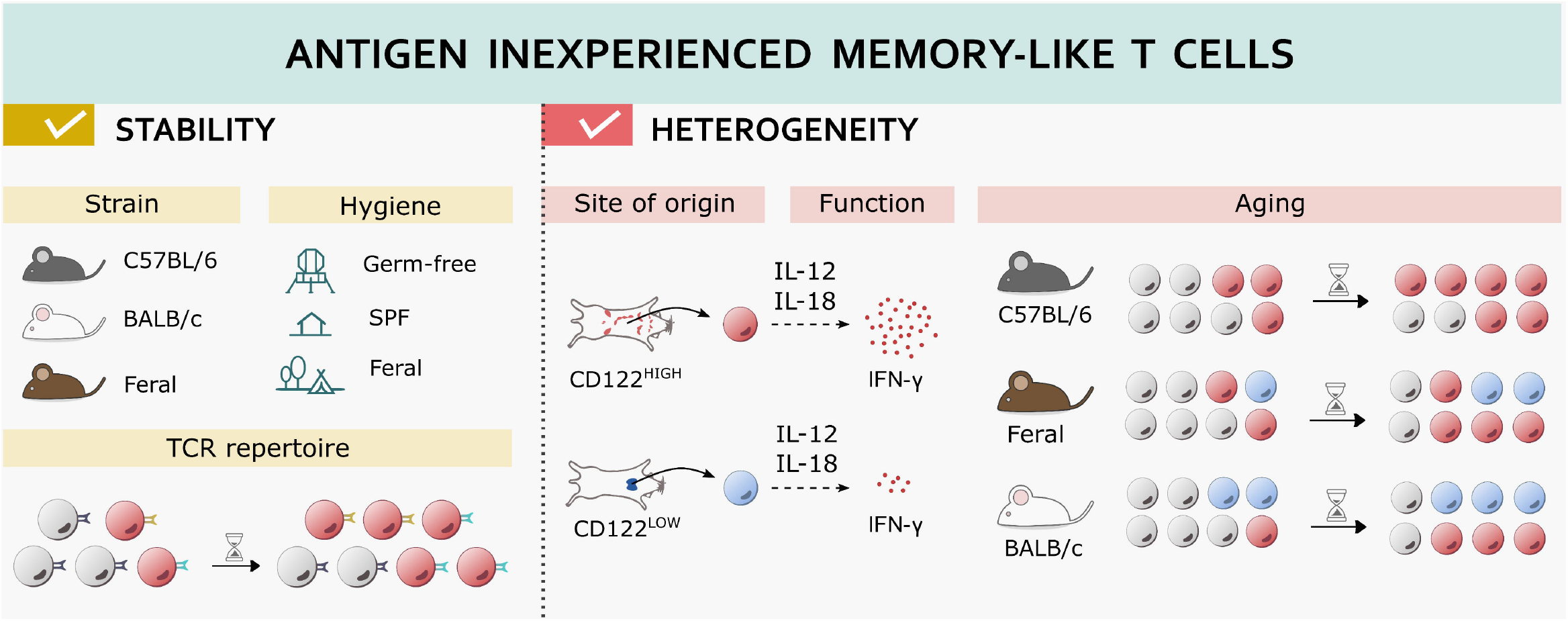

**Figure S1.**
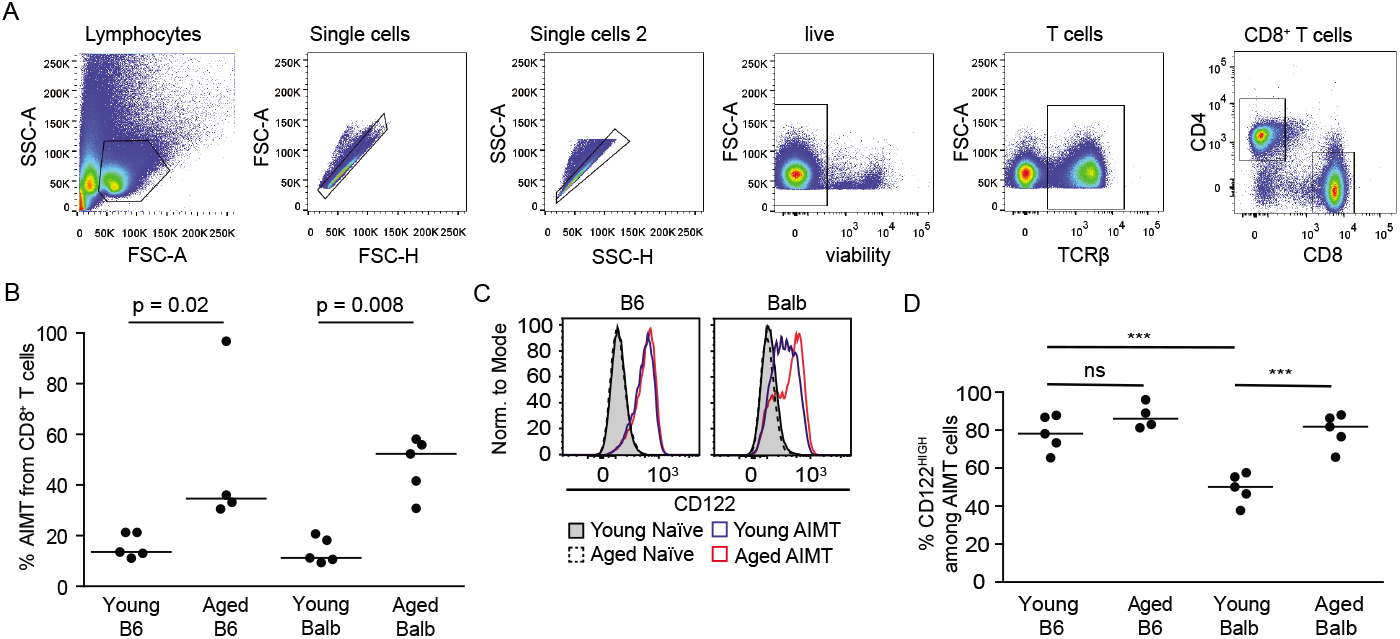

**Figure S2.**
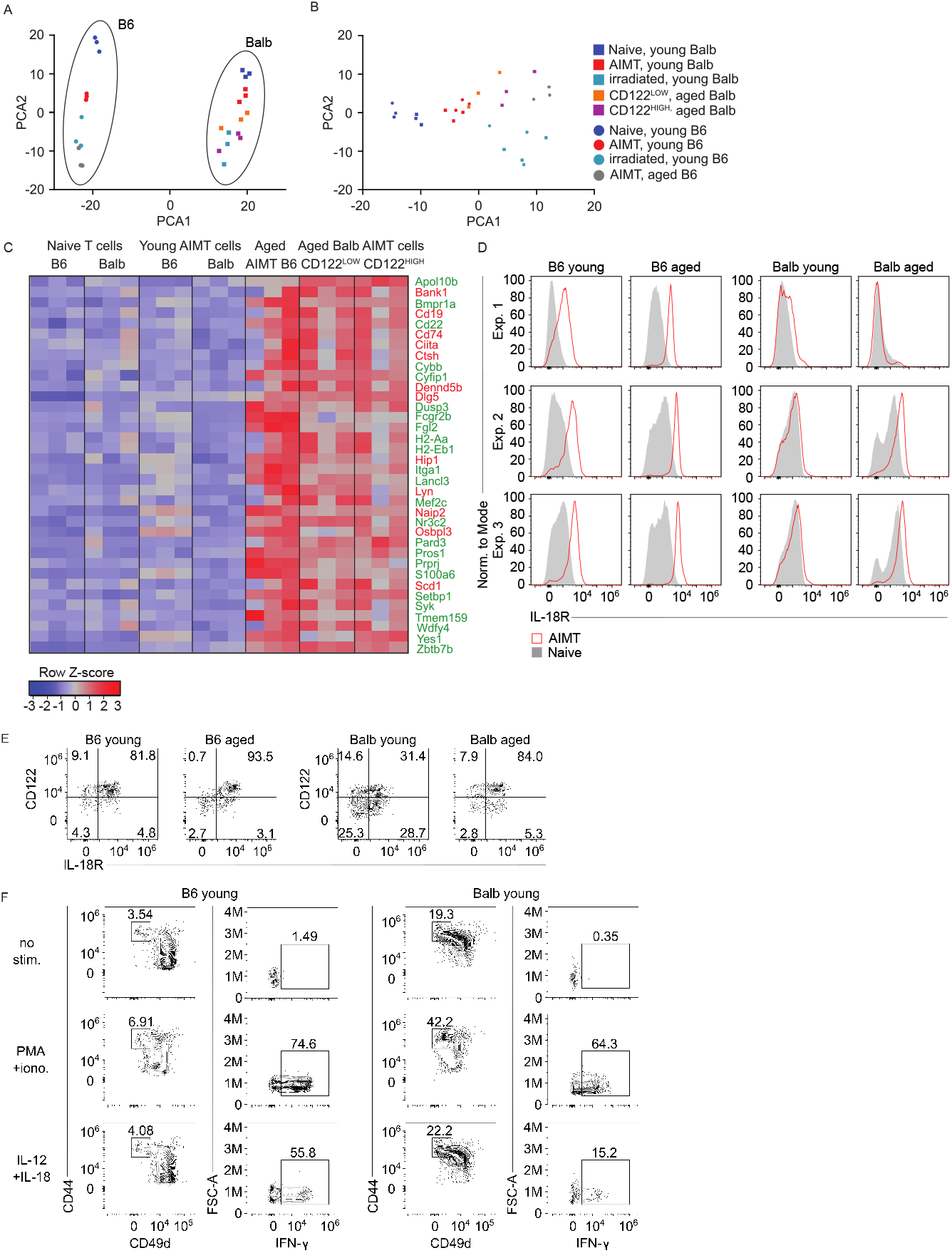

**Figure S3.**
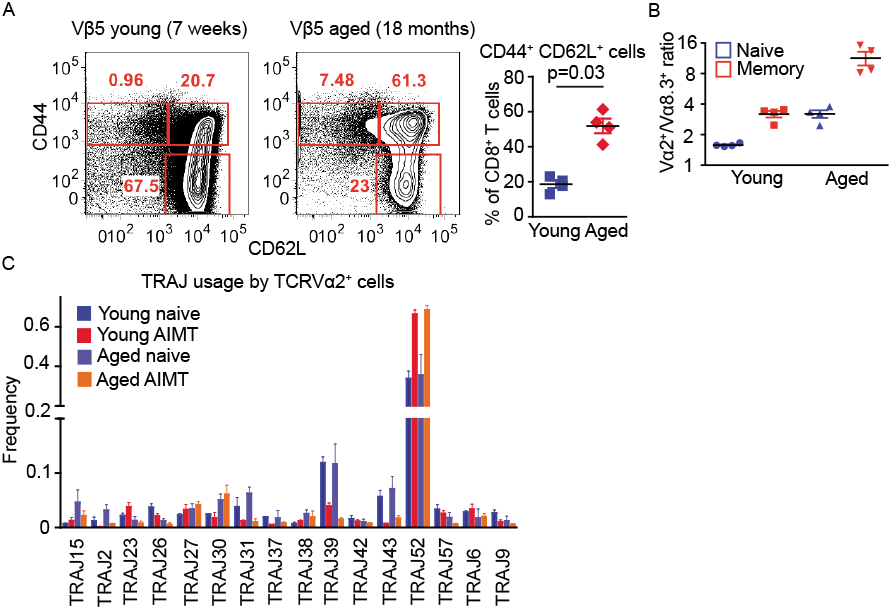

**Figure S4.**
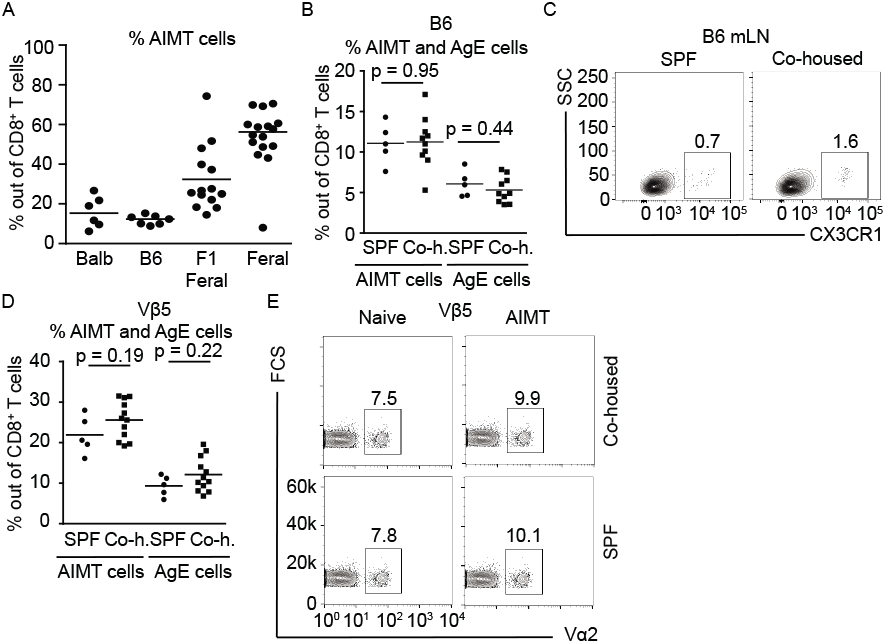

**Figure S5.**
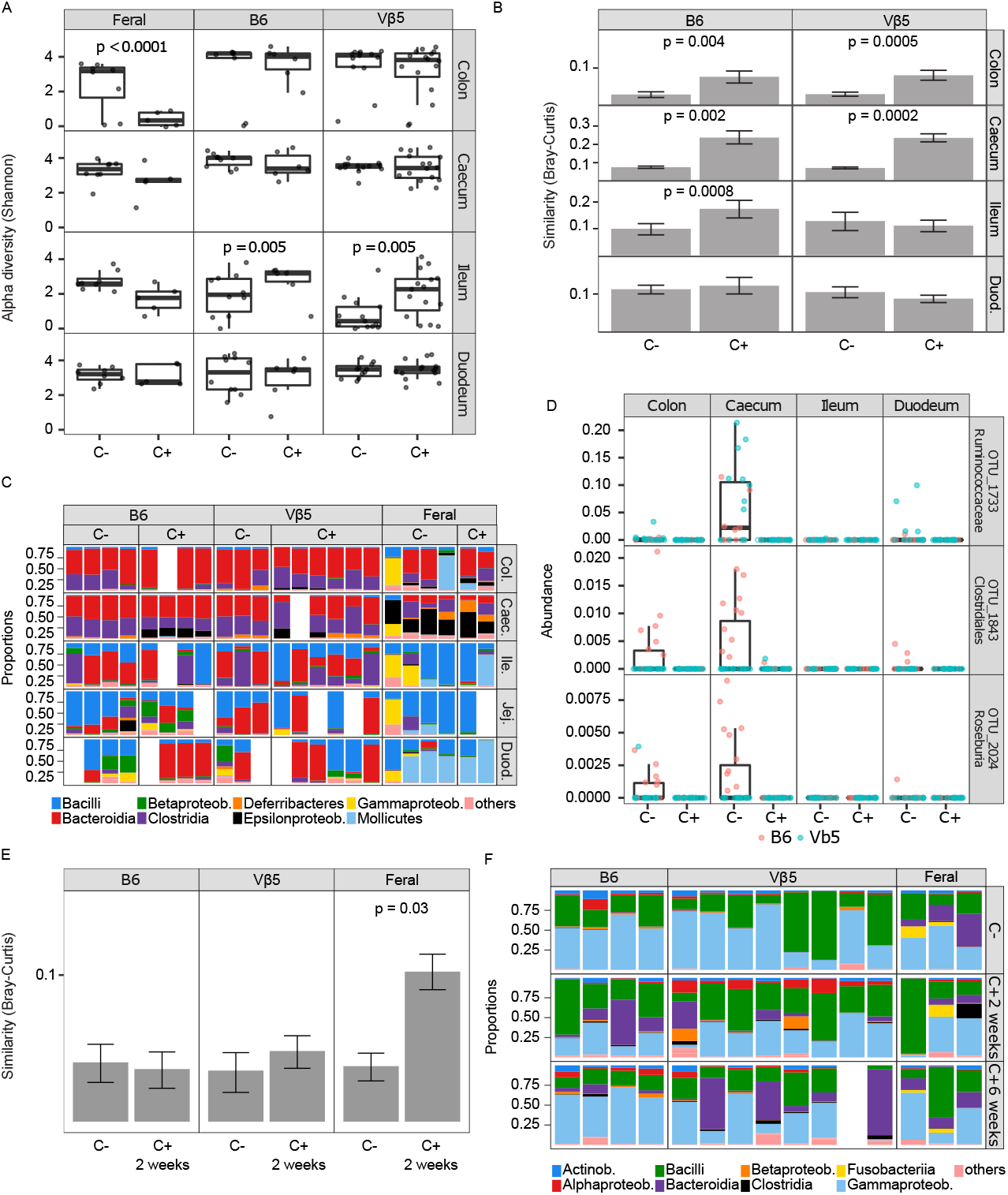

