## Supplementary material for "Phenotypic and clonal stability of antigen-inexperienced memory-like T cells across the genetic background, hygienic status, and aging": Fig. S

**Figure S1**

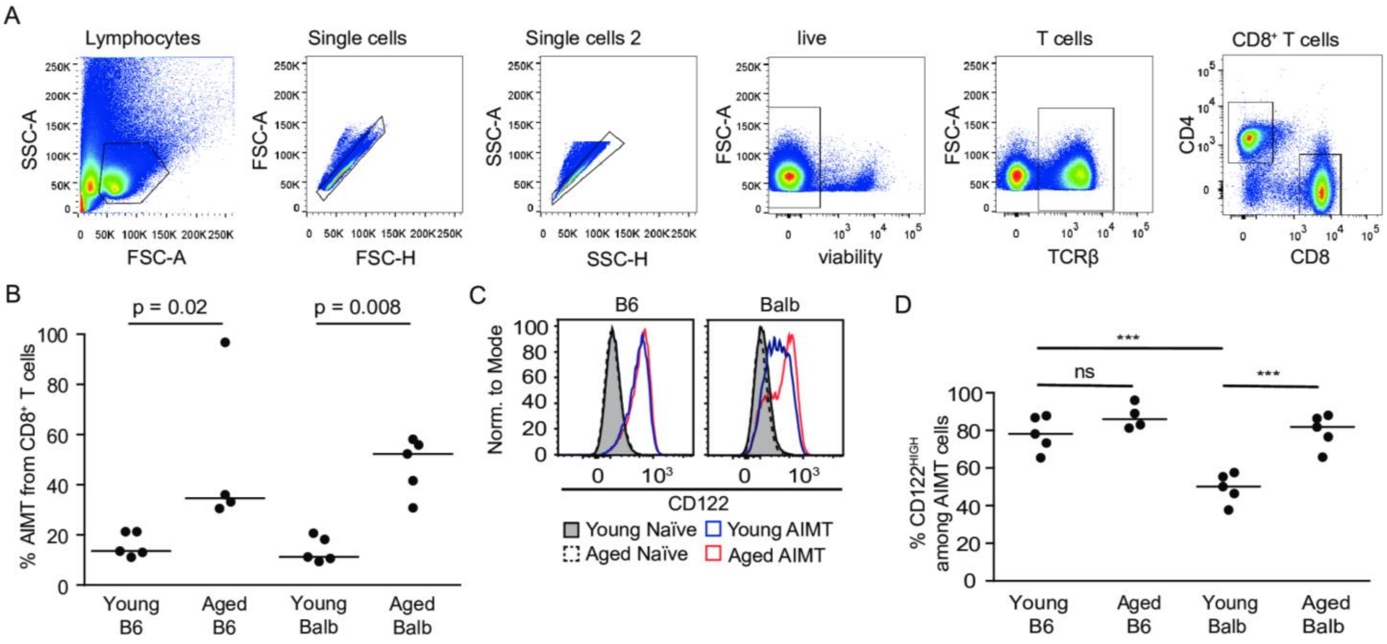


**Supplemental Figure 1. Related to Figure 1**

(A) Gating strategy for gating CD8$^{+}$ T cells in the flow cytometry experiments shown in this study.

(B-D) Analysis of lymph nodes from the same mice as in the experiments in Figure 1A-D. Median.

(B) Quantification of the percentage of CD44$^{+}$ CD49d$^{-}$ AIMT cells among CD8 T cells. n=4 (Balb) or 5 (B6) mice from 2-5 independent experiments.

(C) Histograms of CD122 expression in CD44$^{-}$ naïve and CD44$^{+}$ CD49d$^{-}$ AIMT cells from indicated mice. A representative experiment out of 2 in total.

(D) Quantification of CD122$^{HIGH}$ cells among CD8$^{+}$ AIMT cells. Median. The statistical significance was tested using 1-way ANOVA ($p < 0.0001$) with Bonferroni's Multiple Comparison (post)Tests. *** $p \leq 0.001$.

**Figure S2**

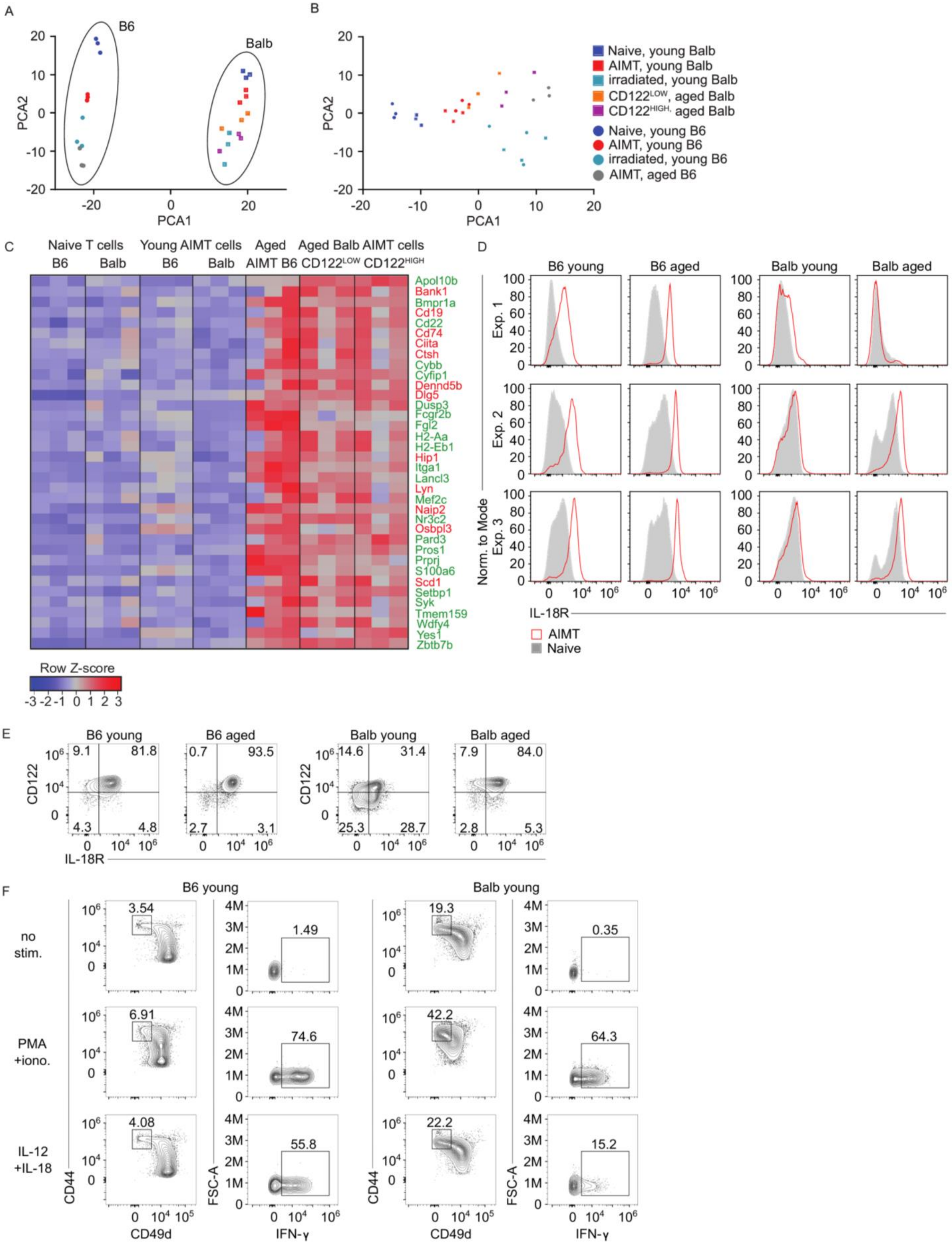

A
B6
Balb
PCA2
PCA1
B
Naive, young Balb
AIMT, young Balb
irradiated, young Balb
CD122LOW, aged Balb
CD122HIGH, aged Balb
Naive, young B6
AIMT, young B6
irradiated, young B6
AIMT, aged B6
C
Naive T cells
Young AIMT cells
Aged AIMT B6
Aged Balb AIMT cells
CD122LOW
CD122HIGH
Apol10b
Bank1
Bmpr1a
Cd19
Cd22
Cd74
Ciita
Ctsh
Cybb
Cyfip1
Dennd5b
Dlg5
Dusp3
Fcgr2b
Fgl2
H2-Aa
H2-Eb1
Hip1
Itga1
Lancl3
Lyn
Mef2c
Naip2
Nr3c2
Osbpl3
Pard3
Pros1
Prprj
S100a6
Scd1
Setbp1
Syk
Tmem159
Wdfy4
Yes1
Zbtb7b
Row Z-score
D
B6 young
B6 aged
Balb young
Balb aged
Exp. 1
Exp. 2
Exp. 3
Norm. to Mode
IL-18R
AIMT
Naive
E
CD122
IL-18R
9.1
81.8
4.3
4.8
0.7
93.5
2.7
3.1
14.6
31.4
25.3
28.7
7.9
84.0
2.8
5.3
F
no stim.
PMA +iono.
IL-12 +IL-18
3.54
1.49
19.3
0.35
6.91
74.6
42.2
64.3
4.08
55.8
22.2
15.2
CD44
CD49d
FSC-A
IFN-γ

**Supplemental Figure 2. Related to Figure 3**

(A-B) PCA analysis (top 500 variable genes) of the gene expression profiles of the individual samples (see Fig. 3) prior to the normalization between strains (A) and after removing genes differentially expressed between strains (B).

(C) Heatmap showing relative expression of 36 genes showing significant upregulation in AIMT cells from aged mice in comparison to AIMT cells in young mice in B6 and Balb strains (both in $CD122^{HIGH}$ and $CD122^{LOW}$ AIMT cells from aged Balb mice). Names of genes upregulated in AIMT cells from aged B6 mice in comparison to young B6 mice in the previously published dataset [36] are in green.

(D) Surface levels of IL-18R in naïve and AIMT $CD8^+$ T cells from young and aged Balb and B6 mice measured by flow cytometry. Supplemental histograms for the experiment shown in Fig. 3G. Three independent experiments are shown.

(E) Surface levels of IL-18R and CD122 in AIMT $CD8^+$ T cells from young and aged Balb and B6 mice measured by flow cytometry. A representative experiment out of three in total (same experiments as shown in Figure S2E).

(F) Production of IFN-γ by AIMT cells (gated as $CD8^+$ $CD44^+$ $CD49d^-$) isolated from young B6 or Balb mice measured by flow cytometry. Supplemental dot plots for the experiment shown in Fig. 3H. A representative experiment out of three in total.

**Figure S3**

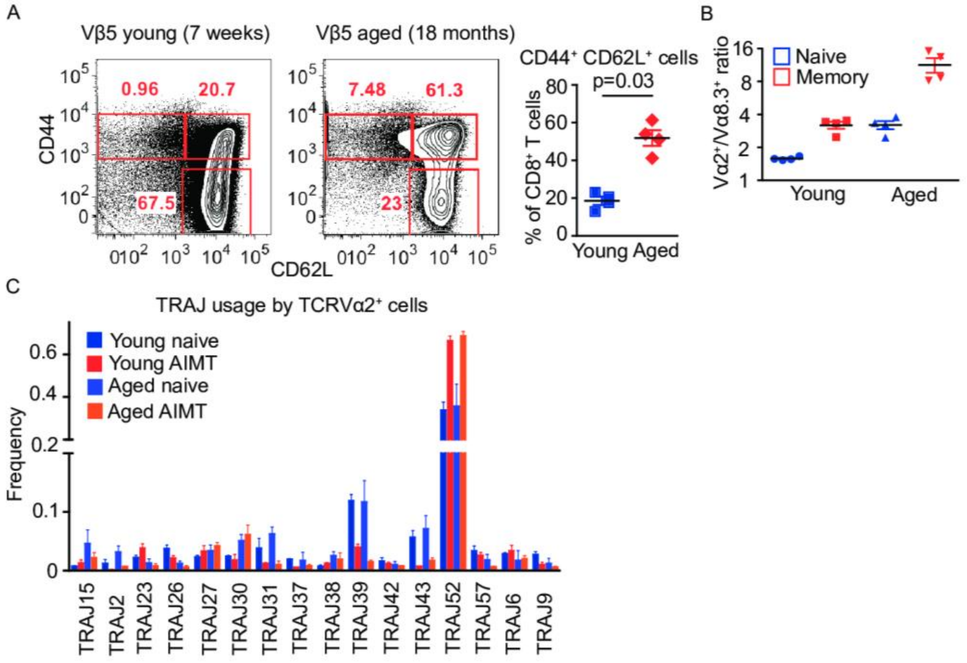


**Supplemental Figure 3. Related to Figure 4.**

(A-B) Analysis of the same young and aged Vβ5 mice as the experiment in Fig. 4A.

(A) Percentage of naïve ($CD44^-$ $CD62L^+$), central memory ($CD44^+$ $CD62L^+$), and effector/memory ($CD44^+$ $CD62L^-$) cells in the lymph nodes of young and aged mice is shown. A representative experiment and the quantification of 4 mice per group are shown. Statistical significance was calculated using Mann-Whitney test.

(B) Quantification of the ratio between the TCRVα2 and TCRVα8.3 T cells among AIMT or naïve $CD8^+$ T cells in Vβ5 young or aged mice.

(C) Quantification of the TRAJ usage by the indicated mice. Mean + SEM. Analysis of the same experiment as shown in Fig. 4B-F.

### Figure S4

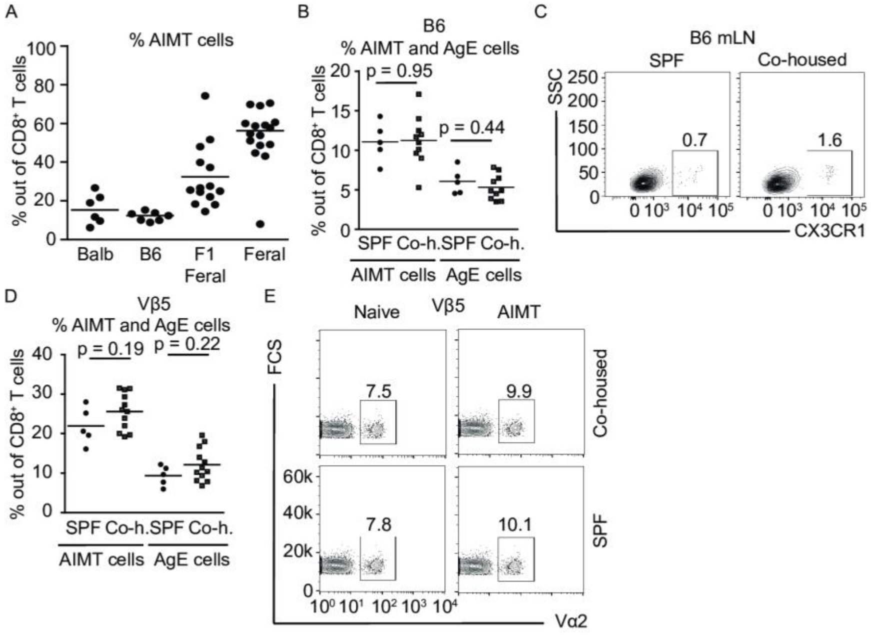


**Supplemental Figure 4. Related to Figure 5.**

(A) Quantification of the percentage of AIMT cells in spleens of Balb, B6, F1 offspring of feral mice, and feral mice. Balb: n = 6 mice/6 independent experiments, B6: 7/7, F1 offspring of feral mice: 14/5, feral mice 16/2. The data are from the same experiments as shown in Fig. 5A-B.

(B) Percentage of $CD44^+$ $CD49d^-$ AIMT and $CD44^+$ $CD49d^+$ antigen experienced (AgE) cells among $CD8^+$ T cells in spleens of SPF B6 mice and B6 mice co-housed with feral mice. The same experiment as shown in Figure 5C-D. Statistical significance was calculated using Mann-Whitney test.

(C) Percentage of $CX3CR1^+$ in $CD8^+$ T cells in spleens of SFP and co-housed B6 mice. The same experiment as shown in Fig. 5C-D.

(D) Percentage of $CD44^+$ $CD49d^-$ AIMT and $CD44^+$ $CD49d^+$ antigen experienced (AgE) cells among $CD8^+$ T cells in spleens of SPF Vβ5 mice and Vβ5 mice co-housed with feral mice. The same experiment as shown in Fig. 5E. Statistical significance was calculated using Mann-Whitney test.

(E) A representative experiment showing the percentage of TCRV$\alpha 2^+$ T cells among AIMT ($CD44^+$ $CD49d^-$) and naïve ($CD44^-$) in SPF and co-housed Vβ5 mice. The same experiment as shown in Fig. 5F.

### Figure S5

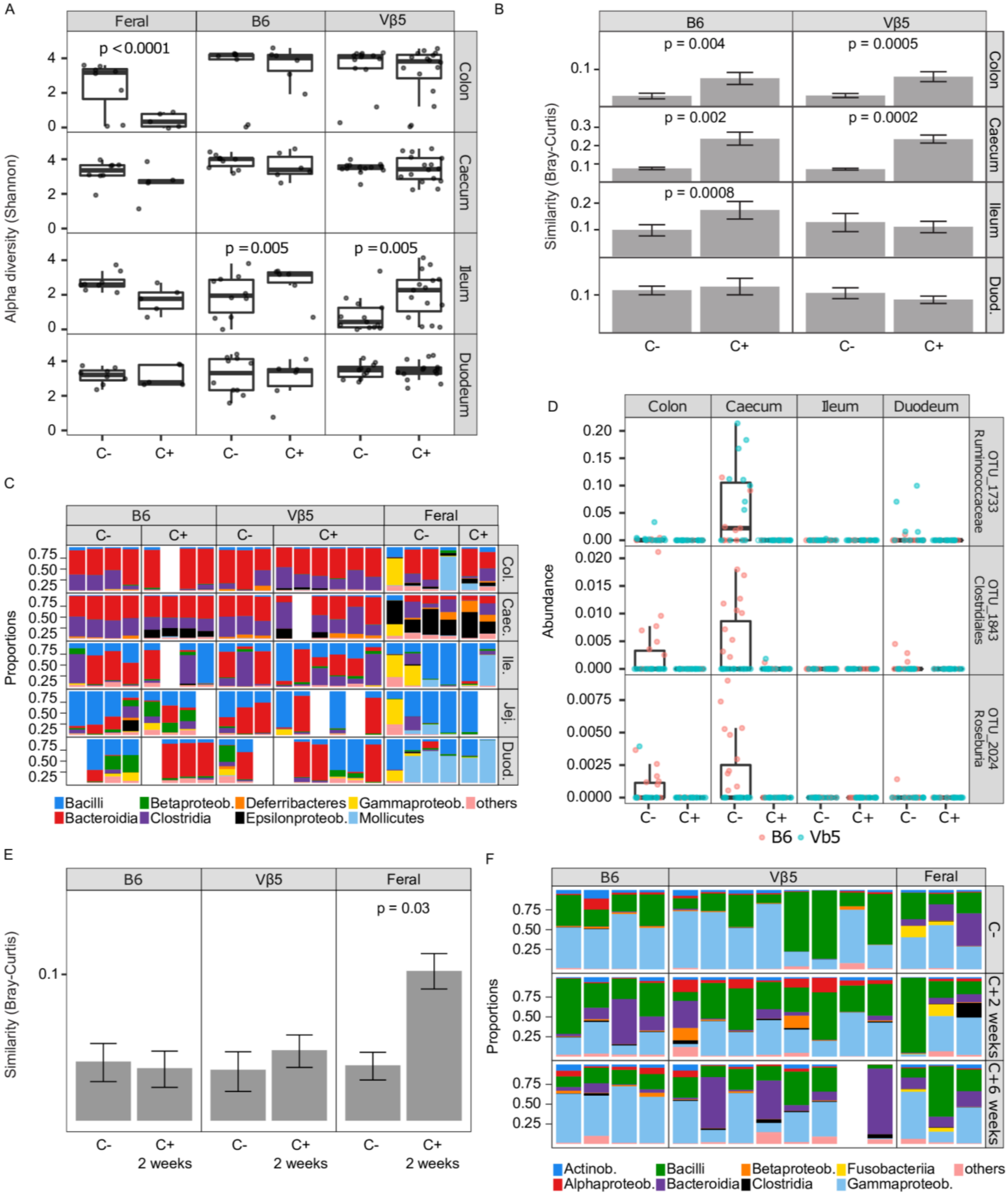
A
Feral
B6
Vβ5
p < 0.0001
p = 0.005
p = 0.005
Colon
Caecum
Ileum
Duodeum
Alpha diversity (Shannon)
C-
C+
B
B6
Vβ5
p = 0.004
p = 0.0005
p = 0.002
p = 0.0002
p = 0.0008
Similarity (Bray-Curtis)
Colon
Caecum
Ileum
Duod.
C
Proportions
Col.
Caec.
Ile.
Jej.
Duod.
Bacilli
Betaproteob.
Deferribacteres
Gammaproteob.
others
Bacteroidia
Clostridia
Epsilonproteob.
Mollicutes
D
Colon
Caecum
Ileum
Duodeum
Abundance
OTU_1733 Ruminococcaceae
OTU_1843 Clostridiales
OTU_2024 Roseburia
B6
Vb5
E
Feral
p = 0.03
Similarity (Bray-Curtis)
C+ 2 weeks
F
Proportions
C+2 weeks
C+6 weeks
Actinob.
Bacilli
Betaproteob.
Fusobacteriia
others
Alphaproteob.
Bacteroidia
Clostridia
Gammaproteob.

**Supplemental Figure 5. Related to Figure 6.**

(A-B) Analysis of the experiment shown in Figure 6A-C.

(A) Shannon-index based alpha diversity of the gut microbiota of feral and laboratory mice co-housed together (C+) or non-co-housed (C-).

(B) Average Bray-Curtis similarity score of intestinal microbiota between laboratory B6 or Vβ5 mice co-housed or non-co-housed with feral mice and non-co-housed feral mice. Error bars correspond to 95% bootstrap confidence intervals, permutation-based p-values are shown for significant differences ($p < 0.05$).

(C) Taxonomical composition of the intestinal microbiota of laboratory co-housed and non-co-housed B6, Vβ5, and feral mice (Experiment A, see Methods). Color bars represent proportions of dominant bacterial classes in each sample.

(D) Three most abundant operational taxonomic units with lower relative abundance in co-housed than non-co-housed B6 and Vβ5 laboratory mice Duod. – duodenum, Ruminococ. – Ruminococcaeae. (Experiment B, see Methods)

(E) Average Bray-Curtis similarity of the salivary microbiota of co-housed (C+) or non-co-housed (C-) laboratory B6 or Vβ5 mice with feral mice to conventional feral mice (left, center), or co-housed or non-co-housed feral mice to laboratory B6 and Vβ5 controls (right) in the experiment shown in Figure 6D. Error bars correspond to 95% bootstrap confidence intervals. Permutation-based p-values are shown for significant differences ($p < 0.05$).

(F) Taxonomical composition of the salivary microbiota of laboratory B6 or Vβ5 mice co-housed (C+) vs. non-co-housed (C-) with feral mice (Experiment A, see Methods). Color bars represent proportions of dominant bacterial classes in each sample. Samples were collected from each mouse prior to the co-housing (C-) and after 2 and 6 weeks during the co-housing experiment.
